# Telomere elongation in the gut extends zebrafish lifespan

**DOI:** 10.1101/2022.01.10.475664

**Authors:** Mounir El Maï, Jean-Marie Guigonis, Thierry Pourchet, Da Kang, Jia-Xing Yue, Miguel Godinho Ferreira

## Abstract

Telomere shortening is a hallmark of aging and is counteracted by telomerase. The gut is one of the earliest organs to exhibit short telomeres and tissue dysfunction during normal zebrafish aging. This is recapitulated in prematurely aged telomerase mutants (*tert-/-*). Here, we show that gut-specific telomerase activity in *tert-/-* zebrafish prevents premature aging. Induction of telomerase rescues gut senescence and low cell proliferation to wild-type levels, while restoring gut tissue integrity, inflammation, and age-dependent gut microbiota dysbiosis. Remarkably, averting gut dysfunction results in a systemic beneficial impact. Gut-specific telomerase activity rescues premature aging markers in remote organs, such as the reproductive (testes) and hematopoietic (kidney marrow) systems. Functionally, it also rescues age-dependent loss of male fertility and testes atrophy. Finally, we show that gut-specific telomerase activity increases the lifespan of telomerase mutants. Our work demonstrates that delaying telomere shortening in the gut is sufficient to systemically counteract aging in zebrafish.

## INTRODUCTION

The discovery that lifespan can be genetically extended in *C. elegans*, initiated a new era of research aiming to define interventions that promote lifespan and healthspan extension^1^. Since then, health and lifespan improvements have been achieved by modulating hallmarks of aging and provided promising therapeutical targets for healthy aging^2^. For example, reverting age-related deregulation of nutrient-sensing mechanisms by interventions such as caloric restriction or rapamycin (mTOR inhibitor) treatment was observed to increase lifespan in several species^3,4^. Similarly, genetic or pharmacologic removal of senescent cells can delay age-associated defects resulting in lifespan extension in mice^5,6^.

Telomere shortening and damage are major determinants contributing to aging^2^. Telomeres protect chromosome ends from degradation and recognition by DNA damage response pathways^7^. Due to the “end-replication problem”, telomeres gradually shorten with each round of cell division^7^. When telomeres become critically short, DNA damage responses are triggered and culminate in cell cycle arrest and, eventually, replicative senescence^8,9^. Consequently, reduced proliferation and accumulation of senescent cells result in loss of tissue integrity^2^. Telomere shortening is counteracted by a specific reverse transcriptase termed telomerase. Telomerase is a multi-subunit ribonucleoprotein, with TERT being its main catalytic component. TERT expression is limited mainly to stem or progenitor cells^10,11^. However, telomerase activity is insufficient to prevent telomere attrition during aging^11^.

Telomeropathy patients carry mutations in telomerase or telomere maintenance protein genes, which lead to premature shortening of telomeres and short life expectancy^12,13^. Similarly, telomerase deficiency in *tert-/-* zebrafish accelerates telomere shortening, leading to premature aging phenotypes and reduced lifespan already in the first generation^14–16^. The majority of tissue dysfunction events described during natural zebrafish aging are anticipated during *tert-/-* zebrafish aging^14–16^. The gut is one of the first organs to exhibit DNA damage associated with short telomeres, reduced cell proliferation, senescence and functional defects not only in natural aging but also throughout *tert-/-* life^14,17^. Importantly, telomere shortening accelerates cellular and functional defects in the gut at a time when other organs remain clear of tissue dysfunction^14^. As in zebrafish, the human gastrointestinal system is one of the organs with the fastest rate of telomere shortening^18^. Telomeropathy patients are often associated with gastrointestinal syndromes^19,20^ and increased telomere shortening was observed in the intestinal epithelium of inflammatory bowel disease patients^21,22^. Therefore, gut homeostasis is intricately connected to telomere length.

A crucial role of gut homeostasis has been described for organism health. Loss of gut permeability is involved in several disorders such as inflammatory bowel disease, diabetes, chronic heart failure and even Parkinson disease^4^. Modification of gut microbiota content (dysbiosis) is associated to aging^23,24^ and is involved in age-related systemic inflammation^25^. Even though weakening of the intestinal barrier is as major feature of gut aging^4^, it remains unclear whether gut aging influences overall organismal aging. Inflammatory and SASP (senescence-associated secretory phenotype) factors chronically emanating from intestinal epithelium with critically short telomeres may impact systemic homeostasis.

Considering that the gut is one of the first organs exhibiting telomere-dependent aging, we anticipated that delaying gut aging would be beneficial for the entire organism. Here, we present a novel vertebrate model aimed at investigating the impact of telomere-dependent gut aging on the entire organism. Using a zebrafish line containing a Cre-inducible and gut-specific *tert* transgene, we show that enterocyte-specific telomerase activity in *tert-/-* fish is sufficient to delay gut aging. Counteracting gut aging improves health of the entire organism, reverting gut microbiota dysbiosis and aging phenotypes in the reproductive and hematopoietic system of *tert-/-* zebrafish. Finally, we show that the most relevant systemic effect of gut-specific telomerase activity is lifespan extension. Thus, gut telomere-dependent aging controls aging of the entire organism.

## RESULTS

### Tissue-specific telomerase activity rescues gut aging

As in humans, the zebrafish gut is one of the organs to exhibit fast telomere length decline^14,18^. To investigate how telomere-dependent gut aging impacts the organism, we generated a Cre-inducible zebrafish transgenic line with gut-specific *tert* expression. This line contains an enterocyte-specific Fabp-2 promoter^26^ upstream of a lox-STOP-lox cassette followed by zebrafish *tert* cDNA in an *tert+/-* genetic background (Figure 1A). After crossing this line with *tert+/-* fish, we induced the *tert* transgene expression by micro-injection of *Cre* mRNA in one-cell stage embryos. Mock injected fish were used as controls for injection and transgene genomic position effects. This experimental set up provided sibling fish that were either *tert-/-* containing the full construct (“*tert-/-* No Cre”; from mock injected embryos), *tert-/-* expressing *tert* transgene (“*tert-/-* +Cre”, from *Cre* mRNA injected embryos), and *tert+/+* containing the full construct (“WT”; from mock injected embryos). Thus, this line allows us to investigate the effects of telomerase activity specifically in the gut of telomerase deficient fish.

**Figure 1:**
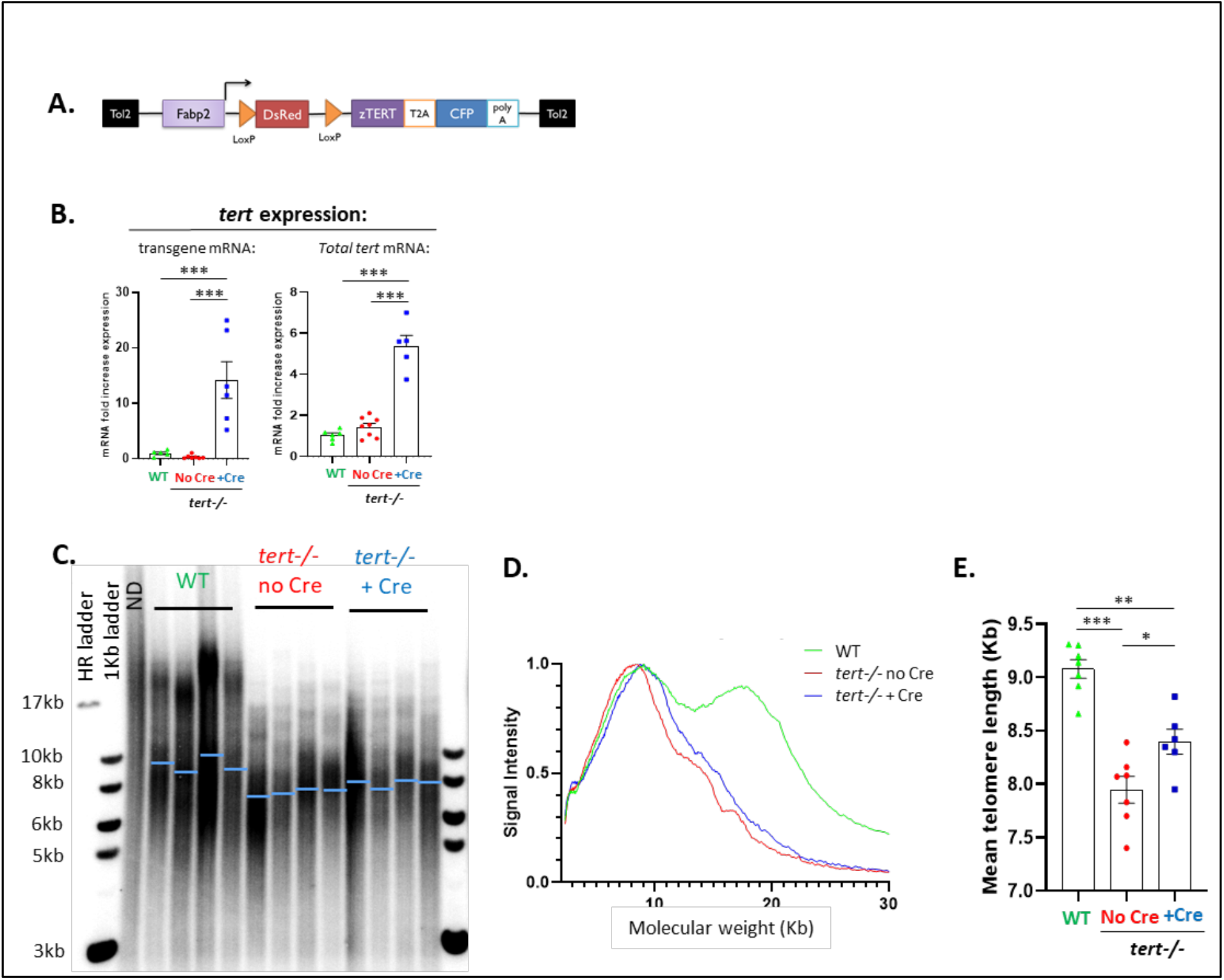
Cre-mediated *tert* expression extend telomere length in *tert-/-* gut tissue. **A**. Schematic representation of the transgene allowing for a Cre inducible and enterocyte specific expression of *tert* mRNA. We created a zebrafish line containing an enterocyte-specific promoter (*Fabp2: fatty acid binding protein 2*) controlling DsRed gene expression flanked by two LoxP sites. Tert-T2A-CFP polycistronic gene was added downstream of the second LoxP site. **B**. RT-qPCR analysis of *tert* transgene mRNA and total *tert* mRNA (endogenous + transgene) expression in 9-month-old gut extracts. RT-qPCR graphs are representing mean ± SEM mRNA fold increase after normalization by *rps11* gene expression levels (N=5-8; ^***^ p-value<0.001, using one-way ANOVA and post-hoc Tuckey tests). Cre mRNA injection at one cell-stage embryos induces the transcription of *tert* transgene mRNA. **C**. Representative images of telomere restriction fragment (TRF) analysis by Southern Blot of genomic DNA extracted from 9-month-old gut samples and quantifications of mean telomere length (blue bars). **D**. TRF mean densitometry curves (N= 6-7). **E**. Quantification of mean telomere length analyzed by TRF. Cre-mediated and enterocyte specific tert expression elongated telomere length in gut of tert-/-fish. Data are represented as mean +/-SEM (N=6-7; ^*^ p-value<0.05; ^**^ p-value<0.01, ^***^ p-value<0.001, using one-way ANOVA and post-hoc Tuckey tests).

While we did not detect expression in mock injected fish, Cre-mediated removal of the STOP cassette triggered the transcription of *tert* transgene in gut tissue (Figure 1B; left panel). This led to ∼5-fold enrichment of total *tert* mRNA (endogenous and transgene *tert* mRNA) in the gut of *tert-/-* +Cre fish when compared to mock injected control tissues (*tert-/-* No Cre and WT) (Figure 1B; right panel). To test whether expression of the *tert* transgene is sufficient to prevent telomere shortening, we performed Telomere Restriction Fragment (TRF) analysis on gut samples of 9-month-old fish. As previously shown, we observed that the range of telomere length in the gut of WT fish exhibits a bimodal pattern (Figure 1C-D)^14,15^. This pattern reflects the differences in telomere length between cell types. Telomere length of WT blood cells is longer (∼19 kb) than other tissues (∼9 kb) leading to a densitometry pattern with two peaks^14,15^. Reflecting the requirement of telomerase activity to sustain long telomeres in blood cells, telomere length of *tert-/-* blood cells is drastically reduced compared to WT (as seen by the loss of the longer telomere peak, Figure 1D)^14,15^. Consequently, *tert-/-* No Cre presented a unimodal TRF pattern in the intestinal tissue. Even though expression of *tert* cDNA driven by the Fabp-2 promoter did not fully restore telomere length to WT levels, induction of the *tert* transgene is sufficient to elongate telomeres in whole gut tissues of *tert-/-* +Cre fish (7.9 kb to 8.4 kb, N=6-7; p<0.05; Figure 1E). Like *tert-/-* No Cre fish, *tert-/-* +Cre fish lacked the higher molecular weight telomere peak, indicating that the *tert* transgene is not expressed in blood cells.

Described as a hallmark of aging, telomere erosion has been proposed as a “molecular clock” defining the number of cell divisions before cell cycle arrest, cell death or replicative senescence^2^. Reduced cell proliferation and accumulation of senescent cells limits homeostatic regeneration and, consequently, causes loss of tissue integrity. In agreement, accelerated telomere shortening in *tert-/-* fish results in premature aging phenotypes^14^. In order to test whether *tert* transgene expression in the gut of *tert-/-* fish rescues local aging defects, we analysed the gut of 9-month-old fish. As previously reported, compared to WT fish, the gut of *tert-/-* No Cre fish showed a reduced proliferation rate^14,15^. Notably, enterocyte-specific telomerase activity rescued the proliferative capacity of this organ to WT levels (Figure 2A). Similarly, SA--galactosidase assays and transcription levels of the senescence-associated genes *p15/16* and *p21* revealed that telomerase activity reduces cell senescence to WT levels (Figure 2B-D). We previously described a cell fate switch from apoptosis to senescence in old *tert-/-* where senescence becomes predominant^17^. At that age, onset of apoptosis becomes indistinguishable between WT and *tert-/-*. Consistently, we detected no difference in number of apoptotic cells in the intestinal epithelium at 9 months of age between WT, *tert-/-* No Cre and *tert-/-* +Cre fish (Supplementary figure 1A).

**Figure 2:**
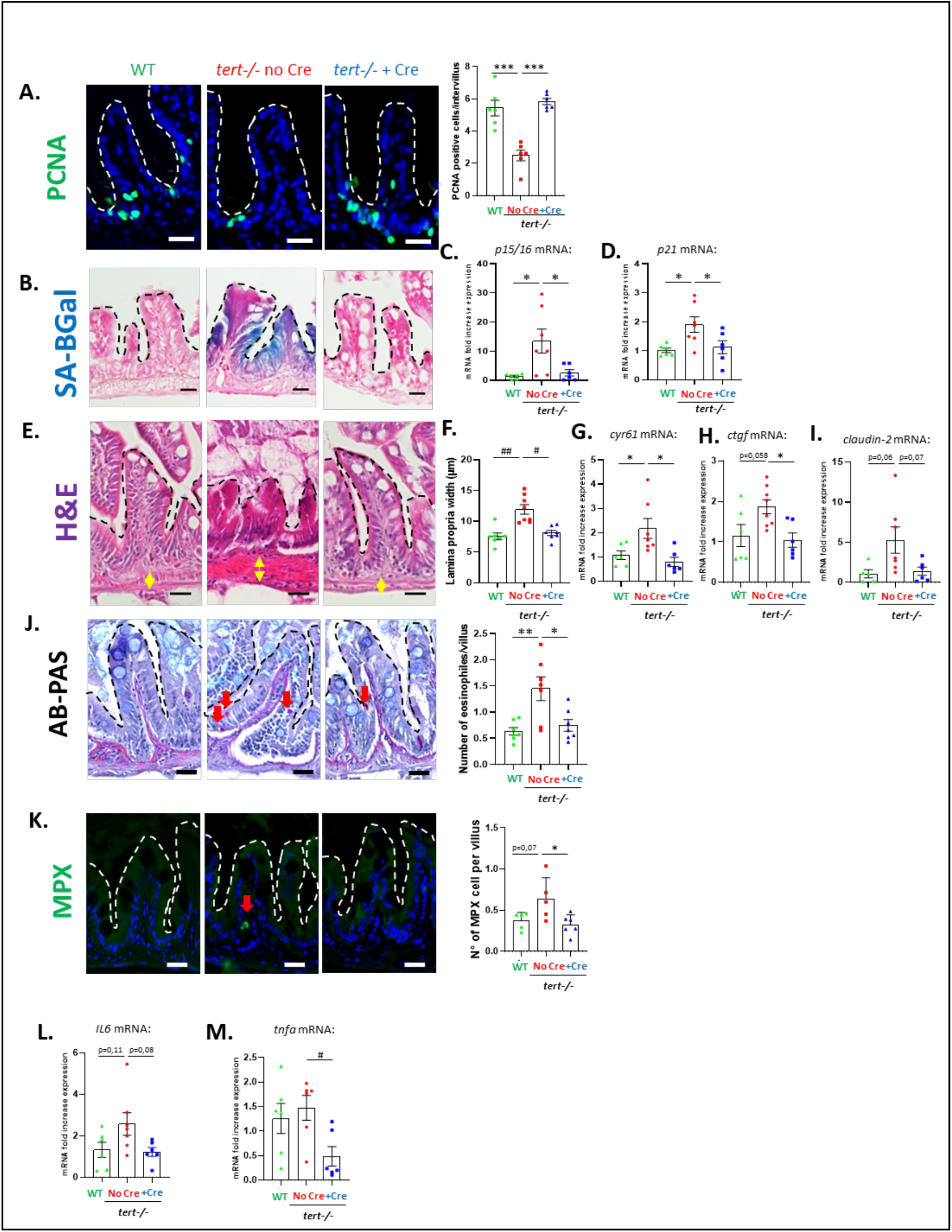
Telomerase reactivation rescues gut aging phenotypes. **A**. Representative immunofluorescence images of proliferation staining (PCNA marker; left panel) and quantification (right panel) in 9-month-old gut tissues. **B**. Representative image of SA--Gal staining of 9-month-old gut cryosection. **C-D**. RT-qPCR analysis of senescence-associated genes *p15/16* (**C**.) and *p21* (**D**.) expression in 9-month-old gut samples. Telomere elongation in gut of *tert-/-* +Cre fish rescues both proliferation and senescence to WT levels compared to *tert-/-* No Cre fish. **E**. Representative hematoxylin and eosin-stained sections of gut from 9-month-old fish (yellow arrows delineate lamina propria width quantified in **F**.). **F**. Quantification of lamina propria width measured on histology images of 9-month-old fish gut. **G-H**. RT-qPCR analysis of YAP target genes *cyr61* (**G.)** and *ctgf* (**H**.) expression in 9-month-old gut samples. **I**. RT-qPCR analysis of the junction protein associated gene *claudin-2* expression in 9-month-old gut samples. **J**. Representative AB-PAS staining images of 9-month-old fish gut (left panel). Number of pink-staining eosinophile cells (red arrows) are quantified in the right panel. **K**. Representative immunofluorescence images of neutrophil staining (MPX marker; left panel) and quantification (right panel) in 9-month-old gut tissues. **L-M**. RT-qPCR analysis of inflammation-associated genes *il6* (**C**.) and *tnfa* (**D**.) expression in 9-month-old gut samples. Telomere elongation in gut of *tert-/-* +Cre fish rescues gut integrity and consequent gut inflammation to WT levels compared to *tert-/-* No Cre fish. Scale bar: 20µm. Dashed lines delineate gut villi. All data are represented as mean +/-SEM (N=6-8 per condition; ^*^ p-value<0.05; ^**^ p-value<0.01, ^***^ p-value<0.001, using one-way ANOVA and post-hoc Tuckey tests; # p-value<0.05; ## p-value<0.01, ### p-value<0.001, using Kruskal-Wallis and post-hoc Dunn’s tests). All RT-qPCR graphs are representing mean ± SEM mRNA fold increase after normalization by *rps11* gene expression levels.

These cellular defects observed in *tert-/-* fish impact tissue integrity^14,15,17^. We observed that 9-month-old *tert-/-* No Cre fish exhibit morphological tissue defects with thickening of the *lamina propria* as compared to WT (Figure 2E-F). Loss of intestinal barrier integrity leads to activation of the YAP (Yes-associated protein) transcription factor responsible for tissue regeneration^27,28^. Consistent with loss of gut integrity, expression of the YAP-target genes *cyr61* and *ctgf* are increased in *tert-/-* No Cre fish compared to WT (Figure 2G-H). Likewise, *claudin-2* mRNA levels are higher in *tert-/-* No Cre compared to WT (Figure 2I). Increased gene expression of the tight-junction protein Claudin-2 occurs during primate aging and enhances *in vivo* intestinal permeability^29,30^. Strikingly, all these phenotypes were rescued in *tert-/-* +Cre fish (Figure 2E-I). As a consequence of loss of intestinal integrity, we observed higher inflammation in the intestinal epithelium of the *tert-/-* No Cre fish as compared to WT. We detected an increased infiltration of eosinophiles (Figure 2J) and neutrophils (Figure 2K) in the gut of *tert-/-* No Cre fish. In line with a rescue of intestinal integrity, the number of these myeloid immune cells was similar to WT in *tert-/-* +Cre fish. Although not significant, gut-specific telomerase activity also ameliorates increased *il6* gene expression observed in *tert-/-* No Cre compared to WT (Figure 2L). Similarly, while no difference was detected in *tnfa* mRNA levels between *tert-/-* No Cre and WT, the expression of this inflammation-related gene is reduced in *tert-/-* +Cre fish (Figure 2M).

By comparing the expression profiles of whole gut tissues using RNA sequencing, we observed a distinguishable transcriptomic signature of *tert-/-* No Cre, while WT and *tert-/-* +Cre samples clustered together (Supplementary figure 2A). GO term analyses showed that, compared to *tert-/-* No Cre, both WT and *tert-/-* +Cre are enriched in gene expression related to cell cycle and ATP production in addition to reduced transcription of genes related to morphogenesis (Supplementary figure 2B-E). Accordingly, KEGG GSEA analyses showed an increase in ribosome and oxidative phosphorylation in these two groups compared to *tert-/-* No Cre, and a decrease of phagosome, cytokine signalling and neuroactive ligand receptor interaction which encompasses arachidonic inflammatory pathway (Supplementary figure 2F-G). In line with the previous results, these transcription profiles confirmed that telomerase activity rescued cell proliferation defects, loss of tissue integrity and inflammation seen in gut of *tert-/-* No Cre fish.

### Local effects: Gut-specific telomerase activity rescues gut microbiota dysbiosis

Gut microbiota dysbiosis is associated with a dysfunctional intestinal barrier and is proposed to generate a feed-forward loop involving gut permeability, inflammation and dysbiosis in aging^25,31^. However, it remains unclear whether delaying gut aging counteracts gut microbiota (GM) dysbiosis. To investigate if telomerase activity in the gut of *tert-/-* fish ameliorates gut dysbiosis, we performed high-throughput sequencing of the V3-4 region of 16S-rDNA of 9-month-old zebrafish gut. Similar to what is described for human aging^23,32^, we observed diminished microbial diversity in *tert-/-* No Cre when compared to WT controls. Both alpha (within samples) and beta (within groups) analyses showed lower diversity in *tert-/-* No Cre individuals compared to WT and *tert-/-* +Cre fish (Figure 3A-B). According to a reduced beta-diversity, using principal coordinates analysis (PCoA), we observed a clustering of *tert-/-* No Cre samples while WT and *tert-/-* +Cre samples were more dispersed (Figure 3C).

**Figure 3:**
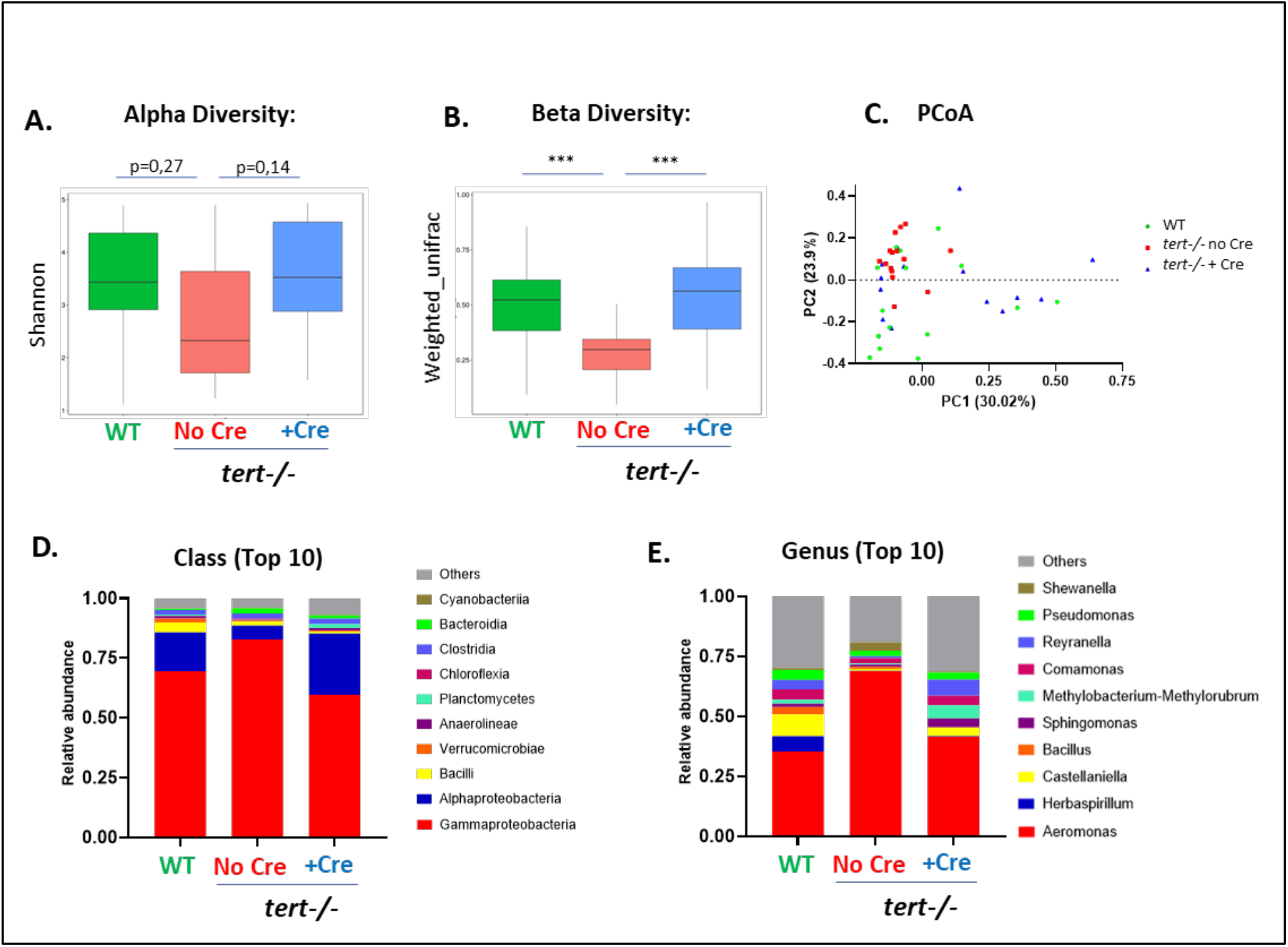
Gut specific telomerase activity rescues gut microbiota dysbiosis. **A**. Quantification of microbiome alpha diversity (within samples) using Shannon index (N=14-15; p-values were determine using Wilcoxon test) in the gut of 9-month-old fish. **B**. Quantification of microbiome beta diversity using weighed unifrac distance (within groups; N=14-15; ^***^ p<0.001 using Tuckey test) in the gut of 9-month-old fish. **C**. Principal Coordinate Analysis (PCoA) of the beta diversity distance (weighted unifrac) in the gut of 9-month-old fish (N=14-15). **D**. Relative abundance of top 10 bacteria classes in the microbiome of the 3 different groups in the gut of 9-month-old fish (N=14-15). **E**. Relative abundance of top 10 bacteria genus in the microbiome of the 3 different groups in the gut of 9-month-old fish (N=14-15). Telomere elongation in gut of *tert-/-* +Cre fish rescues gut microbiota composition and diversity to WT levels compared to *tert-/-* No Cre fish which exhibit gut microbiota dysbiosis.

Relative abundance analysis of bacterial taxonomic units (BTUs) at the class level revealed an overall alteration of GM composition in *tert-/-* No Cre fish compared to WT that was recovered by *tert* transgene expression (Figure 3D). At the class level, we observed in the *tert-/-* No Cre group a decreased abundance of Alpha-proteobacteria and Planctomycetes along with an enrichment in Gamma-proteobacteria, Bacteroidia and Fibrobacteria when compared to other groups (Figure 3D, Supplementary figure 3A). Interestingly, while Alpha-proteobacteria are known to inhibit host cell death and promote proliferation^33^, Gamma-proteobacteria expansion is associated with early age-dependent loss of intestinal barrier integrity in flies^31^.

Similarly, at the genus level, Alpha-proteobacteria *Reyranella* and *Defluviimonas* were reduced while Gamma-proteobacteria *Aeromonas* and *Shewanella* along with *Bacteroides*, a Bacteroidia-related genus, were enriched in *tert-/-* No Cre fish when compared to other groups (Figure 3E; Supplementary figure 3B). Both *Shewanella* and *Aeromonas* genus were described as deleterious in human, with *Shewanella* causing intra-abdominal infections^34^, and *Aeromonas* being associated with inflammatory bowel diseases and inflammation^35,36^. Interestingly, within the *Aeromonas* genus, *A. veronii* species were strikingly overrepresented in *tert-/-* No Cre compared to the other groups (Supplementary figure 3C). From the Bacteroidia class, *B. uniformis, P. merdae* and *B. ovatus* were similarly enriched in *tert-/-* No Cre and can be considered as “pathobionts” that profit from a dysregulated environment to overtake commensal symbionts and become pathogenic^37–39^. Overall, the analysis of gut microbiota composition revealed a dysbiotic microbiota in the *tert-/-* No Cre containing less diverse and more pathogenic bacterial community compared to WT that was reverted by gut-specific telomerase activity.

### Local effects: Gut-specific telomerase activity recovers tissue metabolic profile

Metabolomic analyses can determine the physiological and pathological status of a tissue by measuring metabolites representative of intrinsic and extrinsic factors. Changes in metabolism have been associated with aging and might reflect cellular defects, such as gradual mitochondrial dysfunction with age^40,41^. Similarly, we reported that, by 9 months of age, *tert-/-* gut is affected by mitochondrial dysfunction accompanied by lower ATP and high ROS levels^17^. In order to gain insight on the metabolic profile of *tert-/-* No Cre and the extent of metabolic improvement by telomerase activity, we performed a metabolic analysis of whole intestinal extracts.

Clustering analyses on metabolomic profiles revealed that both WT and *tert-/-* +Cre samples clustered tightly while *tert-/-* No Cre samples differed from other groups (Figure 4A-C). Notably, a significant number of metabolites were reduced (621) or enriched (141) in both WT and *tert-/-* +Cre when compared to *tert-/-* No Cre fish (Supplementary figure 4A). Consistent with our previous work^17^, we observed a drastic reduction of energetic metabolites in *tert-/-* No Cre such as ATP, ADP, NADH, NADPH and CoA compared to the other groups (Figure 4D). Following the anaerobic glycolysis pathway, we noticed lower levels of glucose-6-phosphate and fructose-1,6-bisphosphate and higher amounts of pyruvate and lactate (Supplementary figure 4B). Considering that glucose did not vary greatly between the groups, our results suggest that the gut of *tert-/-* No Cre acquired higher levels of anaerobic glycolysis. We also detected higher pentose shunt activity in *tert-/-* No Cre gut, evidenced by increased amounts of ribose-5-phosphate and eythrose-4-phosphate (Supplementary figure 4C). Interestingly, except for citrate levels, all the detected metabolites of the citric acid cycle were elevated in the *tert-/-* No Cre fish compared to the other genotypes (Supplementary figure 5A). Altogether, the gut energetic metabolism of *tert-/-* No Cre fish appears to be engaged in uncoupled oxidative phosphorylation, consistent with the previously observed damaged mitochondria and higher production of ROS and that, by expressing *tert* transgene in the gut, the metabolic alterations is reverted in the entire tissue.

**Figure 4:**
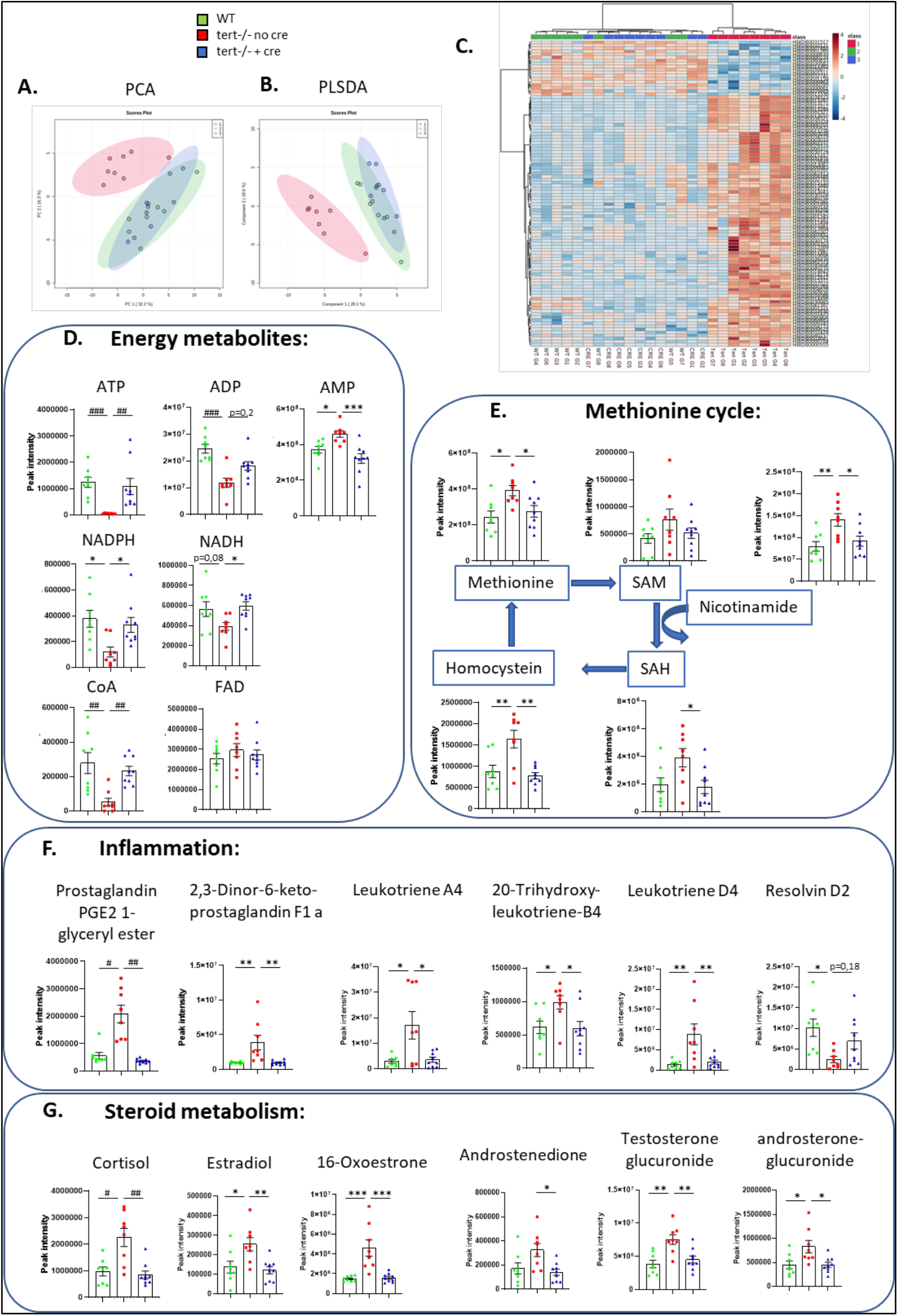
Gut-specific telomerase activity rescues gut metabolomic profile. **A-C**. PCA **(A.)**; Partial Least Squares -Discriminant Analysis (PLSDA)**(B.)**; and metabolite level Heatmap **(C.)** clustering analysis based on untargeted metabolomic data of 9-month-old gut samples. A clustering between *tert-/-* +Cre and WT while *tert-/-* No Cre group was clearly distinguishable from the other (N=8-9 per group). The score plot is represented with a confidence ellipse of 95%. **D-G**. Metabolomic analysis of energy metabolites **(D.)**, methionine cycle pathway **(E.)**, inflammatory metabolites **(F.)** and steroid metabolism **(G.)** in of 9-month-old gut samples. Metabolic alteration seen in gut of *tert-/-* No Cre fish are reverted to WT profile in *tert-/-* +Cre fish. All data are represented as mean +/-SEM (N=8-9 per condition; ^*^ p-value<0.05; ^**^ p-value<0.01, ^***^ p-value<0.001, using one-way ANOVA and post-hoc Tuckey tests; # p-value<0.05; ## p-value<0.01, ### p-value<0.001, using Kruskal-Wallis and post-hoc Dunn’s tests).

Among the detected amino acids, methionine was significantly enriched in *tert-/-* No Cre gut compared to the other genotypes (Figure 4E). We also observed an overall increase in methionine metabolites in the mutant gut that might be allowed by higher levels of nicotinamides.

In line with our previous results depicting higher inflammation of *tert-/-* No Cre fish, we observed an overall increase in the arachidonic metabolism with higher levels of pro-inflammatory molecules, such as prostaglandins and leukotrienes (Figure 4F). Consistently, we detected lower amounts of anti-inflammatory resolvin D2 in *tert-/-* No Cre fish when compared to the other groups. Interestingly, the steroid pathway was also enriched in *tert-/-* No Cre fish. Not only the stress hormone cortisol but also female hormones (such as 16-Oxoestrone or Estradiol) were elevated in male *tert-/-* No Cre fish (Figure 4G). Overall, our unbiased metabolomic analysis described an altered metabolism profile in *tert-/-* No Cre that was recovered by gut-specific telomerase activity.

### Systemic effects: Gut-specific telomerase expression rescues male fertility

In light of the alterations of steroid metabolic profile observed in the gut of *tert-/-* No Cre and to explore the systemic impact of a gut-specific telomerase expression, we analysed aging phenotypes in the male reproductive system. As previously described^14,17^, we observed reduced cell proliferation and high senescence in testes of *tert-/-* No Cre fish (Figure 5A-D). Surprisingly, expression of the *tert* transgene specifically in the gut of *tert-/-* mutants led to a recovery of cell proliferation in the testes. Moreover, SA--Gal and p15/16 mRNA levels were reduced to WT levels in *tert-/-* +Cre testes, while p21 mRNA levels remained similar to *tert-/-* No Cre fish. Similar to what we observed in the gut and consistent to our previous results for 9-month-old fish^17^, apoptotic cell number were similar among the three genotypes (Supplementary figure 1B). Therefore, gut-specific telomerase activity rescues both proliferation and senescence in the reproductive system.

**Figure 5:**
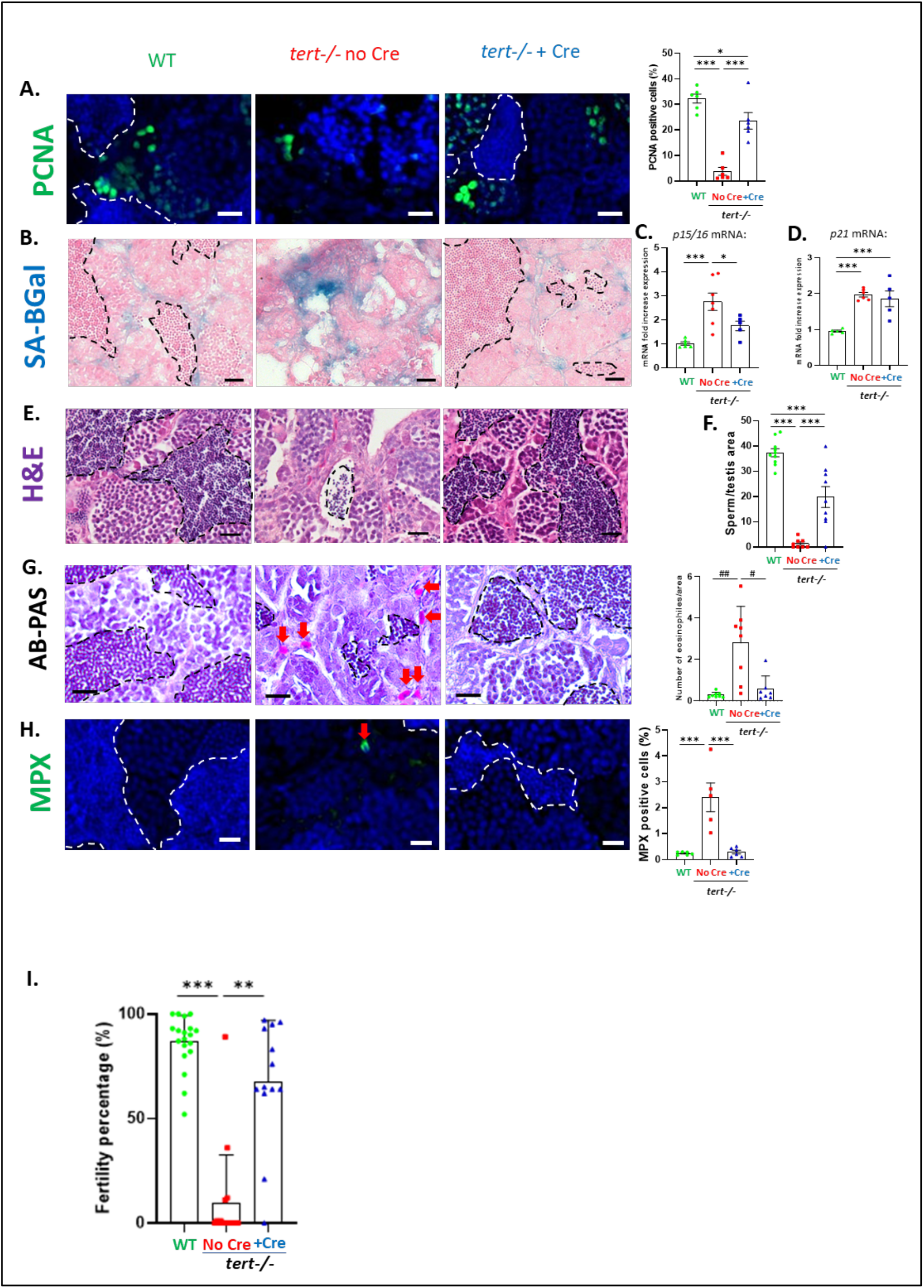
Gut-specific telomerase activity rescues testis aging phenotypes. **A**. Representative immunofluorescence images of proliferation staining (PCNA marker; left panel) and quantification (right panel) in 9-month-old testis tissues. **B**. Representative image of SA--Gal staining of 9-month-old testis cryosection. **C-D**. RT-qPCR analysis of senescence-associated genes *p15/16* (**C**.) and *p21* (**D**.) expression in 9-month-old testis samples. Delaying gut aging in *tert-/-* +Cre fish ameliorates testis proliferation and senescence compared to *tert-/-* No Cre fish. **E**. Representative hematoxylin and eosin-stained sections of testis from 9-month-old fish. **F**. Quantification of mature spermatids area over total testis area measured on histology images of 9-month-old fish testis. **G**. Representative AB-PAS staining images of 9-month-old fish testis (left panel). Number of pink-staining eosinophile cells (red arrows) are quantified in the right panel. **H**. Representative immunofluorescence images of neutrophil staining (MPX marker; left panel) and quantification (right panel) in 9-month-old testis tissues. **I**. Quantification of male fertility of 9-month-old fish determined by counting the percentage of fertilized eggs (detected by successful embryogenesis events) after crossing individually 9-month-old males with a young (3-6 month) WT female. Tert mRNA expression in gut of *tert-/-* (*tert-/-* +Cre fish) have beneficial systemic effects by improving testis function and reducing testis inflammation compared to *tert-/-* No Cre fish. Scale bar: 20µm. Dashed lines delineate mature spermatids area. All data are represented as mean +/-SEM (N=6-8 per condition; ^*^ p-value<0.05; ^**^ p-value<0.01, ^***^ p-value<0.001, using one-way ANOVA and post-hoc Tuckey tests; # p-value<0.05; ## p-value<0.01, ### p-value<0.001, using Kruskal-Wallis and post-hoc Dunn’s tests). All RT-qPCR graphs are representing mean ± SEM mRNA fold increase after normalization by *rps11* gene expression levels.

To ensure that these effects were not due to Fabp2 enterocyte promoter expression in other tissues, we performed RT-qPCR experiments on the testes of the three groups studied. While a clear induction of the *tert* transgene was observed in the gut of 9-month-old *tert-/-* +Cre fish compared to *tert-/-* No Cre fish, no expression of the transgene was detected in the testes (Supplementary figure 6A-B). Accordingly, we could not detect any difference on total *tert* mRNA levels when comparing testes of both groups. Consistently, there was no observable telomere elongation in the testes of *tert-/-* +Cre fish when compared to *tert-/-* No Cre (Supplementary figure 6C-E). As expected, in both these *tert-/-* groups, telomere length was similarly shorter when compared to WT fish. These control experiments support the systemic role of gut-specific telomerase activity in *tert-/-* fish.

Histopathology analysis of testes showed atrophy with a drastically reduced mature spermatids’ content in the *tert-/-* No Cre fish compared to WT (Figure 5E-F), similar to what we previously reported^14,17^. In line with the cell proliferation and senescence rescue, gut-specific telomerase activity recovered these morphological defects. As in the gut, the increased neutrophil and eosinophil testes infiltrates present in *tert-/-* No Cre when compared to WT were also reverted in the *tert-/-* +Cre fish (Figure 5G-H).

Finally, male fertility decreases during natural aging of zebrafish and mice. This loss of fertility is accelerated in the murine and fish premature *tert-/-* aging models^14,42^. To provide further functional insight on the effect of gut aging delay on the reproductive function, we performed a male fertility assay where 9-month-old males of the three groups were individually crossed with young WT females. Percentage of eggs spawned by young females that were fertilized were scored as male fertility index. In accordance with a reduction of mature spermatids’ content, at 9 months of age, *tert-/-* No Cre male fish exhibit a drastic reduction of fertility compared to WT fish (Figure 5J). Strikingly, we observed a full recovery of male fertility in the *tert-/-* +Cre fish. Therefore, gut-specific telomerase activity not only improves cellular and morphological defects of the male reproductive system of *tert-/-* fish, but also rescues their age-dependent loss of fertility.

### Systemic effects: Gut-specific telomerase activity improves health and extends lifespan

We then investigated whether the beneficial effects of gut aging delay could be observed beyond the reproductive system. Given the importance of anemia in telomeropathy patients^43,44^, we specifically investigated for improvements in the kidney marrow, the adult hematopoietic organ in zebrafish. Similar to testes and consistent with the increasing anemic profile, we detected lower levels of cell proliferation in the kidney marrow of *tert-/-* No Cre fish when compared to WT fish (Figure 6A). Notably, upon expression of *tert* transgene in the gut, proliferation rate in the kidney marrow was normalized to WT levels. Similarly, the increased senescence of *tert-/-* No Cre was also rescued in the *tert-/-* +Cre fish (Figure 6B-D). Like in the gut and testes, no differences in apoptosis were detected between the three groups (Supplementary figure 1C). We ruled out Fabp2-dependent expression of telomerase in the kidney marrow of *tert-/-* +Cre fish as we were unable to detect neither *tert* transgene expression nor differences in total *tert* mRNA expression when compared to *tert-/-* No Cre fish (Supplementary figure 6A-B). Consistently, a drastic shortening of telomere length was observed in *tert-/-* +Cre kidney marrow at 9 months of age similar to the telomere length of *tert-/-* No Cre fish (Supplementary figure 6E-H). Therefore, as in testes, gut-specific telomerase activity counteracted telomere-dependent cellular defects in the hematopoietic organ of *tert-/-* fish.

**Figure 6:**
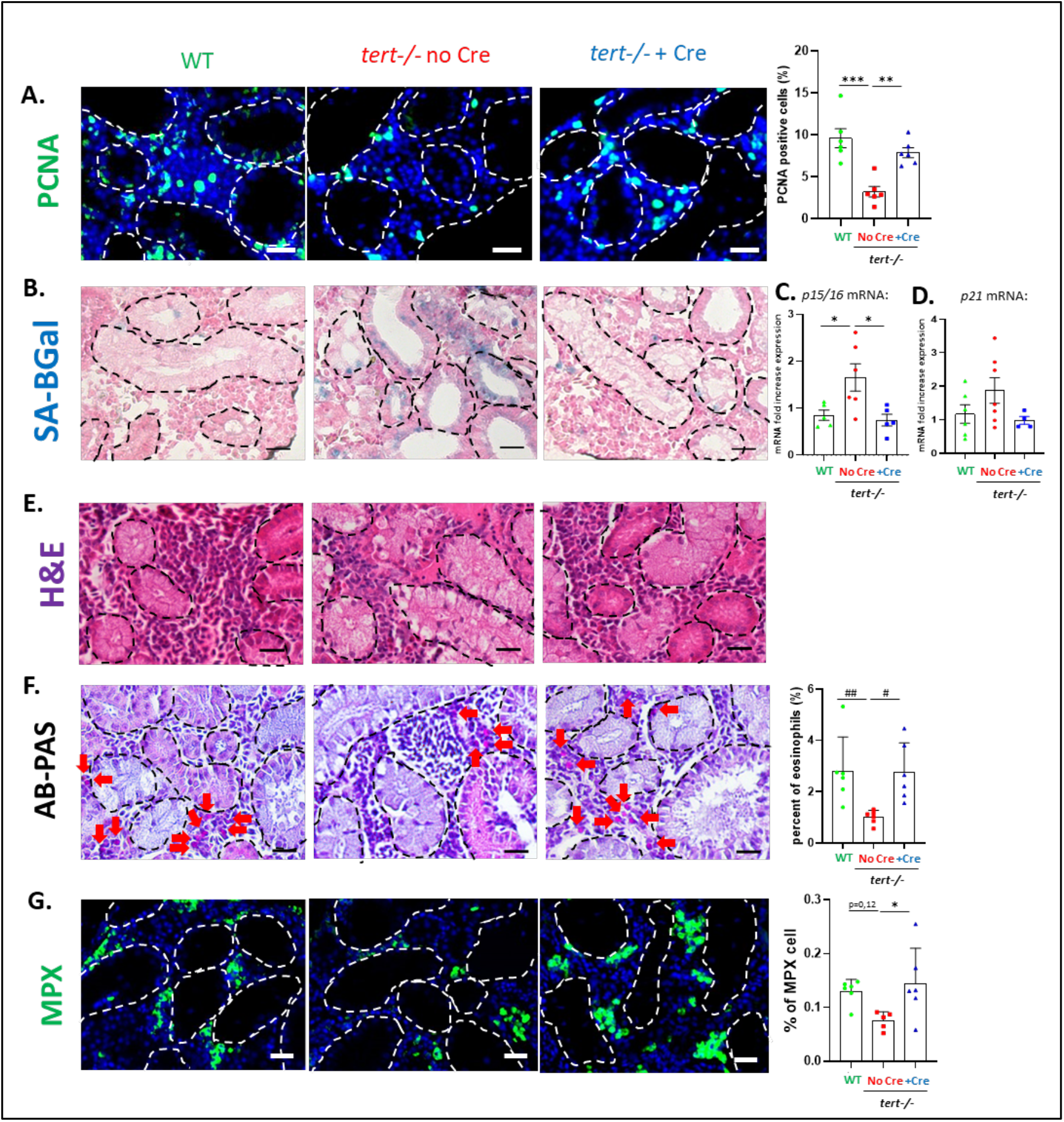
Gut-specific telomerase activity rescues aging of the hematopoietic system (kidney marrow). **A**. Representative immunofluorescence images of proliferation staining (PCNA marker; left panel) and quantification (right panel) in 9-month-old kidney marrow (KM) tissues. **B**. Representative image of SA--Gal staining of 9-month-old KM cryosection. **C-D**. RT-qPCR analysis of senescence-associated genes *p15/16* (**C**.) and *p21* (**D**.) expression in 9-month-old KM samples. Delaying gut aging in *tert-/-* +Cre fish ameliorates KM proliferation and senescence compared to *tert-/-* No Cre fish. **E**. Representative hematoxylin and eosin-stained sections of KM from 9-month-old fish. **F**. Representative AB-PAS staining images of 9-month-old fish KM (left panel). Number of pink-staining eosinophile cells (red arrows) are quantified in the right panel. **G**. Representative immunofluorescence images of neutrophil staining (MPX marker; left panel) and quantification (right panel) in 9-month-old KM tissues. Tert mRNA expression in gut of *tert-/-* (*tert-/-* +Cre fish) have beneficial systemic effects by improving neutrophil and eosinophil pool in compared to *tert-/-* No Cre fish. Scale bar: 20µm. Dashed lines delineate kidney tubules. All data are represented as mean +/-SEM (N=5-8 per condition; ^*^ p-value<0.05; ^**^ p-value<0.01, ^***^p-value<0.001, using one-way ANOVA and post-hoc Tuckey tests; # p-value<0.05; ## p-value<0.01, ### p-value<0.001, using Kruskal-Wallis and post-hoc Dunn’s tests). All RT-qPCR graphs are representing mean ± SEM mRNA fold increase after normalization by *rps11* gene expression levels.

In contrast to other analysed organs, we detected a depletion of immune cells such as eosinophils and neutrophils in the kidney marrow of *tert-/-* No Cre when compared to WT fish (Figure 6F-G). These numbers were reverted to WT levels in *tert-/-* +Cre fish. Our results suggest a decreased reserve pool of eosinophils and neutrophils in *tert-/-* No Cre that is rescued by gut-specific telomerase activity. Decline of immune cells in the kidney marrow may constitute an early sign of hematopoietic dysfunction, comparable to the bone marrow failure described in telomeropathy patients^43,44^.

Finally, considering that delaying gut aging rescued the aging phenotypes of distant organs, such as testes and kidney, we wondered whether telomerase activity in the gut of *tert-/-* would influence zebrafish lifespan. We grew male and female zebrafish of the three different groups and measured their life expectancy. As described previously^14–16^, accelerated telomere shortening of *tert-/-* No Cre fish reduces their lifespan to 12-18 months compared to >42 months in WT fish (Figure 7). Strikingly, delaying gut aging was sufficient to significantly extend the average lifespan of *tert-/-* fish. Average lifespan of *tert-/-* No Cre recovered from 17 months to 24 months in *tert-/-* +Cre fish. However, this was not sufficient to fully rescue life expectancy to WT levels, suggesting that telomere shortening in other organs may become limiting in later stages. Therefore, counteracting gut aging not only delays aging of distant organs, but it is sufficient to extend lifespan of *tert-/-* mutants by 40%.

**Figure 7:**
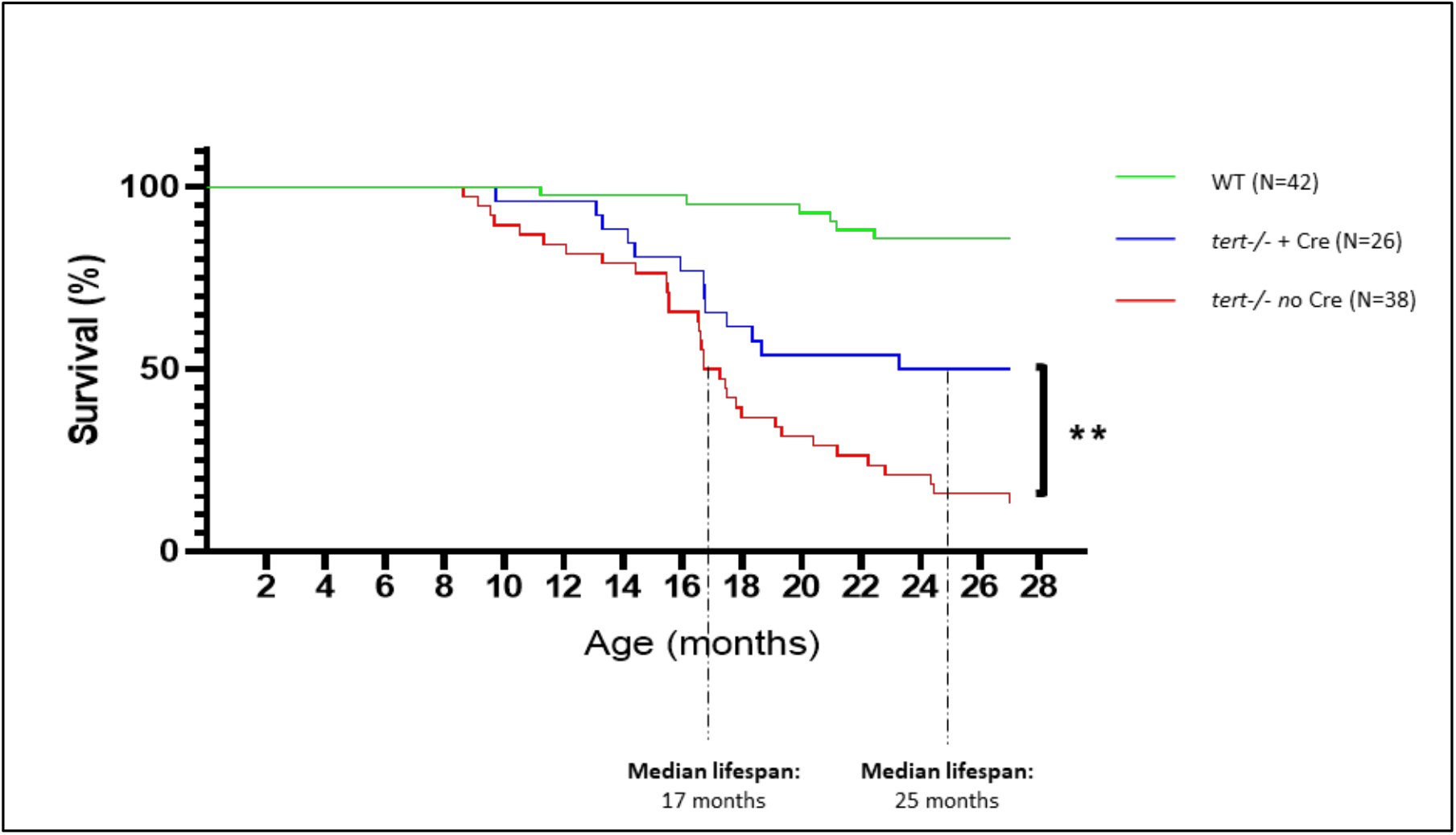
Gut-specific telomerase activity extends lifespan of *tert-/-* zebrafish. Survival curve of WT (N=42), *tert-/-* No Cre (N=38) and *tert-/-* +Cre (N=26) zebrafish. Gut-specific telomerase activity extends lifespan, increasing median life from 17 months in *tert-/-* No Cre to 24 months in *tert-/-* +Cre fish (^**^ p-value<0.01 using Log-rank test).

## DISCUSSION

The gut is a central organ in aging and it constitutes one of the most extensive and selective living barriers to the external environment. Besides its nutrient uptake function, it plays an important role in immune modulation and support a complex interaction with gut microbiota^4^.

In our study, we show that enterocyte-specific telomerase activity in *tert-/-* fish is sufficient to prolong maintenance of gut homeostasis with age. Not only it rescues proliferative defects and cell senescence, but also tissue integrity while reducing tissue inflammation. Interestingly, rescue of gut aging was observed even with a mild but significant telomere extension. Broad telomerase expression counteracts degenerative phenotypes of late generation *tert-/-* mice^45,46^. Improvement of aging phenotypes was observed not only in the gut but also in other organs such testes, spleen, brain or skin. However, in these studies, *tert* expression was not targeted to a specific organ, such as the gut. Consistent with our observations, the dePinho lab recently showed that telomere shortening in the mouse gut activates inflammation by a mechanism involving YAP^47^. In this work, a mosaic expression of *tert* in LGR5 positive cells of *tert-/-* mice partially ameliorated intestinal function and reduced inflammation of this tissue, but no systemic effects were reported apart from body weight rescue and a modest increase in survival. Consistently, we show that YAP target genes were likewise induced in *tert-/-* No Cre fish. These were rescued in the *tert-/-* +Cre fish that, not only reverted the YAP pathway, but also rescued local inflammation. Moreover, we now show that counteracting gut telomere dysfunction also delays remote organ dysfunction and overall organismal aging. It is worth noting that trace amounts of *fabp2* transcripts were previously reported in zebrafish in the liver, brain and kidney marrow but not in testis^48^. While in our study, we did not detect any *fabp2* promoter-dependant transgene expression and telomere extension in kidney marrow, we cannot exclude that a negligible *tert* transgene expression in non-proliferative tissues such as brain and liver might participate to the systemic improvement.

We report that delaying telomere-dependent gut aging has beneficial systemic effects. Expression of *tert* transgene exclusively in the gut of *tert-/-* fish reverted cellular defects in the reproductive (testes) and the hematopoietic (kidney marrow) systems, namely reduced cell proliferation and senescence. At the organ level, improving cellular turnover and reducing inflammation in testes allowed for replenishment of mature spermatids leading to functional rescue of male fertility. In parallel, we observed that neutrophil and eosinophil pools were restored in the hematopoietic system. Strikingly, in line with a systemic recovery, counteracting intestinal aging of *tert-/-* zebrafish extended lifespan by 40%. Notably, our study indicates that proliferative organs such as the reproductive or hematopoietic systems can conserve regenerative capacity even in a context of short telomeres. This has been observed in the rescue of telomerase deficiency by *tp53* mutations in several organisms, namely mice and zebrafish^15,49^. Thus, our study anticipates that maintenance of proliferative capacity and tissue integrity relies on external signals induced by an aging gut.

How would gut aging influence the entire organism? The recent years have seen a flurry of studies supporting the role of inflammation/SASP in inducing paracrine senescence in remote tissues^50,51^. Senescent cells accumulate with age in tissues and promote aging by secreting molecules such as inflammatory cytokines, chemokines and other molecules, also known as SASP^51^. Remarkably, clearance of these cells delay age-associated defects and lead to lifespan extension^5,6^. We previously reported that some organs, such as kidney marrow, exhibit onset of cellular senescence before reaching critically short telomeres during zebrafish aging^14,15^. Interestingly, in the present study, we observed that enterocyte-specific telomerase activity in *tert-/-* fish not only counteracted senescence in the gut but also in distant organs. A paracrine signaling driven by inflammation/SASP factors secreted by an aged gut with short telomeres might therefore promote senescence in remote organs in *tert-/-* No Cre zebrafish. This mechanism would affect cell proliferation systemically and eventually lead to loss of tissue homeostasis in the entire organism.

Over the last decades, gut microbiota has captured the scientific community’s interest by its implications in the etiology of several diseases including inflammatory bowel disease, type 2 diabetes, hypertension, liver diseases and depression^52^. Modification of gut microbiota dysbiosis has been linked to aging^23,24^ and is involved in age-related systemic inflammation^25^. We show that *tert-/-* No Cre fish are afflicted by gut microbiota dysbiosis, with reduced population diversity and enrichment of pathogenic bacteria. Gut-specific telomerase activity recapitulated the bacterial diversity and composition observed in WT fish. Therefore, delaying gut aging counteracted gut microbiota dysbiosis. We anticipate that, as gut telomere shortening becomes limiting, increasingly dysbiotic microbiota exacerbates the age-dependent effects over the entire organism through its microbial components and/or by inducing systemic inflammation. This idea is supported by a recent work showing that stool transfers from young to middle-aged individuals was sufficient to extend lifespan of short-lived killifish^53^.

By analysing the metabolic profile of the gut in our model, we noticed an overall accumulation of methionine and its metabolites (SAM, SAH and homocystein) that is reverted to WT levels in *tert-/-* +Cre fish. Similar enrichment of methionine and its metabolites with age were reported in human and mice^40,54^. Interestingly, dietary methionine restriction or impeding SAM accumulation extends lifespan in different animal models^4,55,56^. Moreover, hyperhomocysteinemia has been implicated in several age-related disorder such as cardiovascular diseases, osteoporosis, renal and cognitive dysfunctions^54^. Mechanistically, deleterious effects of methionine and its metabolites involves DNA methylation drift, mTOR activation, inflammation and oxidative stress^4,41,56^. While it remains unclear why the levels of methionine and its metabolites are enriched in *tert-/-* gut, we anticipate/propose that propagation of these molecules throughout the zebrafish organism may precipitate systemic aging.

A growing list of evidence described the implication of the gut in different systemic physiological and pathological processes, often involving gut microbiota. Overall, the present work describes a central role of telomere shortening in the gut during the aging of a vertebrate organism. While it provides several mechanistic clues, it remains unclear how this organ influences aging of the entire organism. This includes microbiota dysbiosis, inflammation/SASP and dysregulation of methionine metabolism. Finally, our study demonstrates that targeting aging of a unique organ, the gut, is an exciting strategy to extend health span and lifespan.

## MATERIAL AND METHODS

### Ethics statement

Zebrafish work was conducted according to local and international institutional guidelines and were approved in France by the Animal Care Committee of the IRCAN, the regional (CIEPAL Cote d’Azur #697) and national (French Ministry of Research #27673-2020092817202619) authorities and in Portugal by the Ethical Committee of the Instituto Gulbenkian de Ciência and approved by the competent Portuguese authority (Direcção Geral de Alimentação e Veterinária; approval number: 0421/000/000/2015).

### Plasmid construct

Zebrafish *tert cDNA* was obtained using TertFL-pCR-II-Topo plasmid kindly provided by S. Kishi laboratory^57^. Using Gibson assembly recombination methods, *tert* cDNA and *eCFP* cDNA were linked by *T2A* sequence and inserted into *Ubi: loxP-dsRed-loxP-EGFP* vector plasmid (a kind gift from Zon’s lab derived from *ubi:Switch and lmo2:Switch* contructs^58^). Then, *t*he intestine-specific intestinal fatty acid binding protein promoter *(-*2.3kb *fabp2* -also called *ifabp*-) was amplified by high fidelity PCR (iProof_ High-Fidelity DNA Polymerase; Bio-Rad, Hercules, CA, USA) from p5E–2.3ifabp plasmid (kindly gifted by J. Rawls laboratory). -2.3kb fabp2 PCR product was then cloned into the Ubi: LoxP*-dsRed-loxP-tert-T2A-CFP* using sfI/ FseI digestion to provide the final construct: Fabp2: LoxP*-dsRed-loxP-tert-T2A-CFP*.

### Generation of transgenic fish

*Tol2* mRNA was synthesized with SP6 RNA polymerase from pCS2FA-transposase plasmid (Tol2Kit) using mMESSAGE mMACHINE SP6 transcription kit (Invitrogen; Cergy Pontoise, France). One-cell stage zebrafish embryos were micro-injected with 1,4 nL of a mixture containing 25 ng/µL of linearized plasmid and 100 ng/µL of Tol2 mRNA, diluted with RNase-free water. Injected fish were raised to adulthood and germline transmitting fish were then selected and out-crossed to wild type AB until obtaining a single copy transgenic line Tg(Fabp2: LoxP*-dsRed-loxP-tert-T2A-CFP)*.

### Zebrafish lines and maintenance

Zebrafish were maintained in accordance with Institutional and National animal care protocols. Generation and maintenance of the telomerase mutant line *tert* AB/hu3430 (referred in this work as tert+/-) were previously described^14,15,17^. This line was outcrossed with Tg(Fabp2: LoxP*-dsRed-loxP-tert-T2A-CFP)* line to obtain a stock line combining both transgenics. All stocks were kept in heterozygous form for tert mutation and maintained strictly by outcrossing to AB strains to avoid haploinsufficiency effects in the progeny.

Experimental fish were obtained by crossing *tert+/-* fish with *tert+/-;* Fabp2: LoxP*-dsRed-loxP-tert-T2A-CFP*. The generated sibling embryos were then micro-injected with 1.4nL of either 25ng/µL Cre mRNA diluted in RNase-free water (Cre induced fish), or RNase-free water alone (mock injected fish). This experimental set up provided sibling fish that are either tert-/-; Fabp2: LoxP*-dsRed-loxP-tert-T2A-CFP* (mock injected *tert-/-* referred to as *“tert-/-* No Cre”), *tert-/-* ; Fabp2: *tert-T2A-CFP* (Cre induced *tert-/-* referred to as “*tert-/-* +Cre”) or *tert+/+;* Fabp2: LoxP*-dsRed-loxP-tert-T2A-CFP* (mock injected wild type referred to as “WT”). Overall characterization of these three genotypes was performed in F1 sibling animals at 9 months of age. Due to a male sex bias in our crosses, that affected mostly *tert-/-* progeny, we were unable to obtain significant numbers of females for analysis and so all of our data except survival analysis are restricted to males.

### Senescence-associated beta-galactosidase staining

Tissues were fixed with 4% PFA during 3 hours at 4°C. After being washed in PBS, they were incubated in 30% sucrose (Sigma, MO, USA) at 4°C until sinking (24-48hours). Fixed tissues were then embedded in OCT medium (M-M France, Brignais, France) and kept at -80C. Senescence-associated beta-galactosidase staining was then performed on slides of 5µm cryosections using Senescence Beta-Galatosidase staining kit (#9860, Cell Signalling Technology, Danvers, MA, USA) following manufacturer’s instructions. After 16h (testis, kidney marrow) or 3h (gut) incubations with the X-Gal staining solution at 37°C, slides were washed with PBS and counterstained for one minute with Nuclear Fast Red (NFR) solution (Sigma, MO, USA) prior to being dehydrated and mounted.

### Telomere restriction fragment (TRF) analysis by Southern blot

Isolated tissues were first lysed at 50°C overnight in lysis buffer (Fermentas #K0512; Waltham, MA, USA) supplemented with 1 mg/ml Proteinase K (Sigma, MO, USA) and RNase A (1:100 dilution, Sigma, MO, USA). Genomic DNA was then extracted by equilibrated phenol-chloroform (Sigma, MO, USA) and chloroform-isoamyl alcohol extraction (Sigma, MO, USA). Same amounts of gDNA were digested with RSAI and HINFI enzymes (NEB, MA, USA) for 12 h at 37°C. After digestion, samples were loaded on a 0.6% agarose gel, in 0.5% TBE buffer, and run on a CHEF-DRII pulse field electrophoresis apparatus (Bio-Rad). The electrophoresis conditions were as follow: initial swtich 1s, final switch 6s; voltage 4V/cm; at 4°C for 20 h. Gels were then processed for Southern blotting using a 1.6 kb telomere probe, (TTAGGG)n, labelled with [α-32P]-dCTP.

### Fertility assays

In order to assess male fertility, 9-month-old male individuals from the three different genotypes were separately housed overnight in external breeding tanks with a single young (3-6 month old) WT female. Breeding pairs were left to cross and to lay eggs the following morning. Embryos were collected approximately 2 hours post fertilization (hpf) and allowed to develop at 28°C. Assessment of egg fertilization and embryo viability was conducted between 2 and 4 hpf. At least 14 independent crosses were conducted for each genotype to evaluate male fertility. Only successful breeding trials, defined as events where clutch of eggs was laid by a female, were scored.

### Histology

Zebrafish were sacrificed by lethal dose of 1g/L of MS-222 (Sigma, MO, USA), fixed for 72 hr in 10% neutral buffered formalin and decalcified in 0.5M EDTA for 48 hr at room temperature. Whole fish were then paraffin-embedded in order to perform five micrometer sagittal section slides. Slides were stained with haematoxylin and eosin for histopathological analysis. In parallel, slides were stained by Alcian Blue (AB) solution pH 2.5 (Sigma, MO, USA) followed by Periodic acid-Schiff staining (kit #395B, Sigma, MO, USA) according to manufacturer’s instructions. Microphotographs (N>=6 fish per genotype) were acquired in a Leica DM4000B microscope coupled to a Leica DFC425 C camera.

### Immunofluorescence

Deparaffinized and rehydrated slides were microwaved 20min at 550W in citrate buffer (10 mM Sodium Citrate, pH 6) to allow for antigen retrieval. Slides were washed two times in PBS for 5 minutes each and blocked for 1 hour at RT in 0.5% Triton, 5% normal goat serum in PBS (blocking solution). Subsequently, slides were incubated overnight at 4°C with 1:50 dilution of primary antibody in blocking solution. The following primary antibodies were used: mouse monoclonal antibody against Proliferation Cell Nuclear Antigen (PCNA, sc56 Santa Cruz, CA, USA, 1:50 dilution) and rabbit polyclonal against myeloperoxidase (MPX, GTX128379; Irvine, CA, USA, 1:50 dilution). After two PBS washes, overnight incubation at 4°C was performed with 1:500 dilution of goat anti-rabbit or anti-mouse secondary antibodies Alexa Fluor 488 (Invitrogen; Cergy Pontoise, France). Finally, after DAPI staining (Sigma, MO, USA), slides were mounted DAKO Fluorescence Mounting Medium (Sigma, MO, USA). Apoptosis was detected using the In SituCell Death Detection Kit (Roche, Bâle, Switzerland) as previously described^14,17^. Briefly, deparaffinated sections were permeabilized by one hour incubation at 37°C with 40 μg/ml Proteinase K in 10 mM Tris-HCl pH 7.4. After being washed with PBS, slides were incubated one hour at 37°C with TUNEL labelling mix (according to manufacturer’s instructions) prior to DAPI staining and mounting.

Immunofluorescence images were acquired on Delta Vision Elite (GE Healthcare, Chicago, IL, USA) using a OLYMPUS 20x/0,75 objective. For quantitative and comparative imaging, equivalent image acquisition parameters were used. The percentage of positive nuclei was determined by counting a total of 500–1000 cells per slide (N>=6 zebrafish per genotype).

### Real-time quantitative PCR and RNA sequencing

Zebrafish were sacrificed by lethal dose of 1g/L of MS-222 (Sigma, MO, USA) and each tissue (gonads, gut and kidney marrow) were dissected and immediately snap-frozen in liquid nitrogen. RNA extraction was performed by disrupting individual tissues with a pestle in TRIzol (Invitrogen, UK) followed by chloroform extractions. Quality of RNA samples was assessed through BioAnalyzer (Agilent 2100, CA, USA). Retro-transcription into cDNA was performed using QuantiTect Reverse Transcription kit (Qiagen, Hilden, Germany).

Quantitative PCR (qPCR) was performed using FastStart Universal SYBR Green Master mix (Roche, Bâle, Switzerland) and an 7900HT Fast Real-Time PCR Detection System (Thermofisher, Waltham, MA, USA). qPCRs were carried out in triplicate for each cDNA sample. Relative mRNA expression was normalized against *rps11* mRNA expression using the 2^_ΔΔCT^ method as compared to controle condition. Primer sequences are listed in Supplementary table.

RNA sequencing was performed by the Beijing Genomics Institute (BGI; Hongkong), using for each condition, biological triplicates consisting for each of a pool of two individual tissues. DNAse treated total RNA samples were enriched for mRNAs using oligo dT magnetic beads. In turn, mRNAs were fragmented into 200 bp-size fragments and the first strands of cDNAs were synthesized by using random-hexamers. In order to generate library products, double stranded cDNAs from the second strand synthesis were then purified by magnetic beads followed by A-tailing and RNA adaptors ligation. The library was amplified with phi29 to make DNA nanoball (DNB) which had more than 300 copies of one molecular. Pair ended 150 bases reads were sequenced in the way of combinatorial Probe-Anchor Synthesis (cPAS) on DNBseq plateform and 100M clean reads per sample was generated. Raw data with adapter sequences or low-quality sequences was filtered SOAPnuke software developed by BGI. The RNA-seq reads were analyzed via an internal pipeline for transcript quantification, normalization, and comparison. Briefly, the human reference genome assembly vGRCh38 (retrieved from http://www.ensembl.org) and gencode annotation v37 (retrieved from https://www.gencodegenes.org/) were processed by gffread v0.12.2 to extract human reference transcriptome. Based on this extracted reference transcriptome, Salmon v1.4 was used to perform transcript quantification via quasi-mapping. RUVseq v1.20.0 was used for data transformation by “rlog” and data normalization by replicates. DESeq2 v1.26.0 was used for differentially expressed gene (DEG) analysis. The false discovery rate (FDR) cutoffs of 0.1 were explored for the DEG analysis. Based on the resulting DEG candidate gene lists, clusterProfiler v4.0 was employed for Gene Ontology (GO) analysis and Gene Set Enrichment Analysis (GSEA), based on which KEGG pathway enrichment analyses were further performed.

**Supplementary Table.**
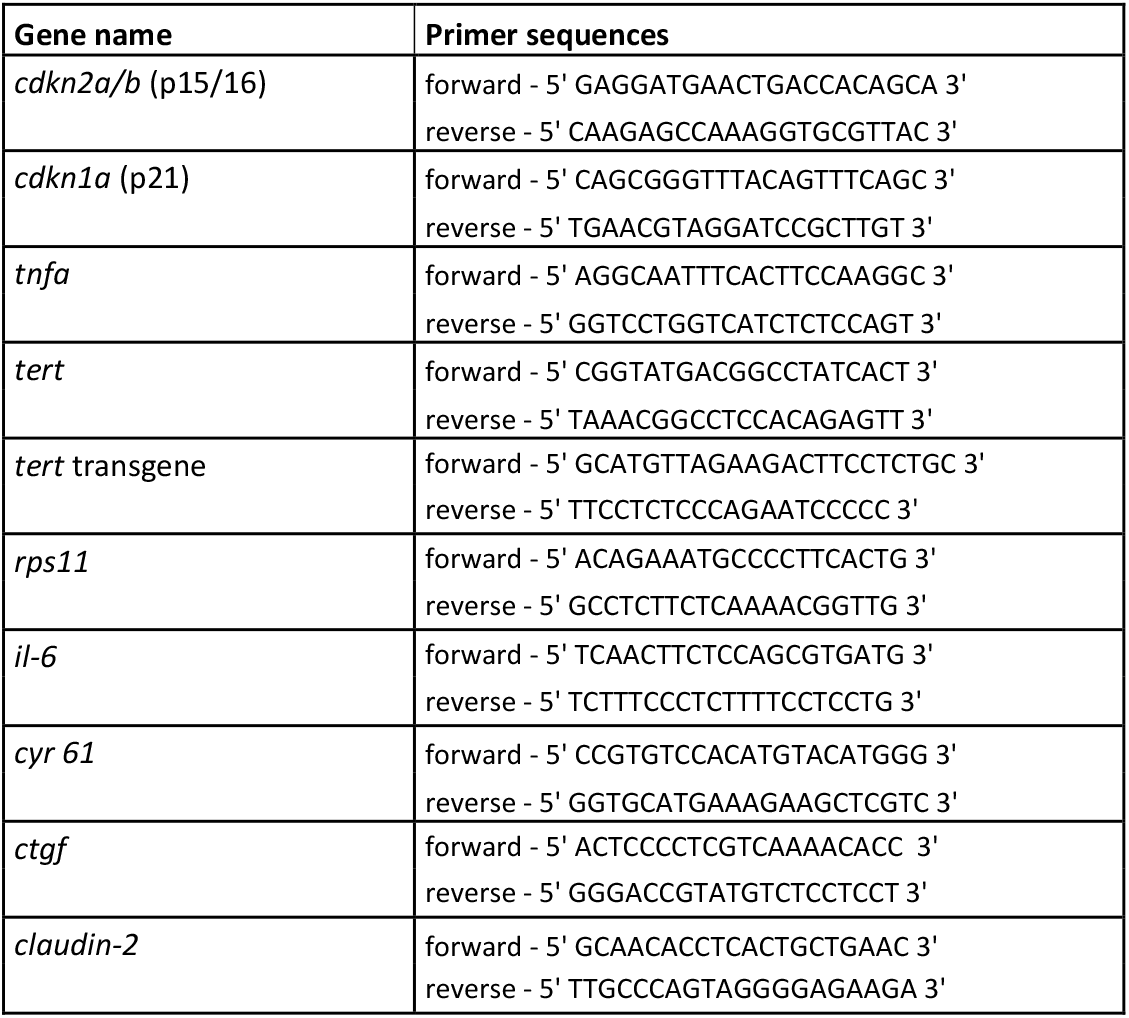
List of primers used in RT-qPCR expression analysis.

### Metagenomics

gDNA was first extracted from isolated gut of sibling fish as described for Telomere restriction fragment (TRF) analysis. The V3-V4 hypervariable regions of bacterial 16S rRNA genes were amplified by PCR with Phusion® High-Fidelity PCR MasterMix (New England Biolabs, Ipswich, MA, USA) using specific primer as previously described^59^. PCR products were mixed at equal density ratios and purified with Qiagen Gel Extraction Kit (Qiagen, Germany). The sequencing libraries were generated using NEBNext® UltraTM DNA Library Prep Kit and sequenced on Illumina NovaSeq 6000 paired-end platform to generate 250 bp paired-end raw reads. Sequences analysis were performed using Uparse software with all the effective tags. Sequences with ≥97% similarity were assigned to the same OTUs. Representative sequence for each OTU was screened for further annotation. For each representative sequence, Mothur software was performed against the SSUrRNA database of SILVA Database for species annotation at each taxonomic rank (Threshold:0.8∼1). QIIME and R were used to calculate alpha and beta diversity metrics and generate plots. Principal Coordinate Analysis (PCoA) was performed to get principal coordinates and visualize from complex, multidimensional data.

### Metabolomic analysis

Each frozen gut sample was homogenized in methanol 600 µL of methanol (HPLC grade, Merck Millipore, USA) and incubated overnight at -20°C. Tubes were vortexed and incubated overnight at - 20°C for protein precipitation. After centrifugations, supernatants were removed, dried using a SpeedVAC concentrator (SVC100H, SAVANT, Thermo Fisher Scientific, Illkirch, France), resuspended in 80 µL of a 20:80 acetonitrile-H2O mixture (HPLC grade, Merck Millipore) and stored at -20°C until use for metabolomic analysis.

Chromatographic analysis was performed with the DIONEX Ultimate 3000 HPLC system coupled to a chromatographic column (Phenomenex Synergi 4 u Hydro-RP 80A 250_3.0 mm) set at 40°C and a flow rate of 0.9 mL/min. Gradients of mobile phases (mobile phase A: 0.1% formic acid in water and mobile phase B: 0.1% formic acid in acetonitrile) were performed over a total of 25 min. MS analysis was carried out on a Thermo Scientific Exactive Plus Benchtop Orbitrap mass spectrometer. The heated electrospray ionization source (HESI II) was used in positive and negative ion modes. The instrument was operated in full scan mode from m/z 67 to m/z 1000. The post-treatment of data was performed using the MZmine2 version 2.39 (http://mzmine.github.io/). Metabolites were identified using the Human Metabolome Database version 5.0 (http://www.hmdb.ca). We only used ions identified as [M+H]^+^ adducts in the positive mode and [M-H]^-^ adducts in the negative mode and ions found in all the samples after gap filling.

### Statistical analysis

Graphs and statistical analyses were performed in GraphPad Prism8 software (San Diego, CA, USA), using one-way ANOVA test with Tuckey’s post-correction or Kruskal-Wallis test with Dunn’s post-hoc test. A critical value for significance of p<0.05 was used throughout the study. For survival analysis, Log-rank tests were performed using GraphPad Prism8 in order to determine statistical differences of survival curves.

Untargeted metabolomic analysis of gut samples were processed using statistical analysis [one factor] modules proposed by MetaboAnalyst 5.0 (https://www.metaboanalyst.ca). For each comparison, peak intensities were Log transformed. Clustering analysis were performed using Principal Component Analysis (PCA), Partial Least Squares - Discriminant Analysis (PLS-DA), and Heatmap tools provided by MetaboAnalyst.

## ACKNOWLEDGEMENTS

We thank members from the Telomeres and Genome Stability and the Telomere Shortening and Cancer Laboratories for fruitful discussions. We are grateful to Leonor Saúde (Instituto de Medicina Molecular) and Ana Rita Araújo (IPMC) for critically reading our paper. This work was supported by the Fondation Arc pour la Recherche sur le Cancer (PJA20161205137) and the Fondation pour la Recherche Médicale FRM (EQU201903007804). MEM was supported by a postdoctoral fellowship from the Ville de Nice. This work was also supported by the Université Côte d’Azur - Académie 4 (Installation Grant: Action 2 - 2019) and the Howard Hughes Medical Institute International Early Career Scientist grant awarded to MGF. We thank the Instituto Gulbenkian de Ciência (IGC) histology unit, the IGC imaging unit, for assistance with experimental planning, sample processing and data collection and the IGC Fish Facility for excellent animal care. IGC Fish Facility is financed by Congento LISBOA-01–0145-FEDER-022170, co-financed by FCT (Portugal) and Lisboa2020, under the PORTUGAL2020 agreement (European Regional Development Fund). The work was also performed using the PEMAV fish facility, Imaging core facility (PICMI) and the Genomics facilities at the IRCAN supported by FEDER, Région Provence Alpes-Côte d’Azur, Conseil Départemental 06, ITMO Cancer Aviesan (plan cancer), Cancéropole Provence Alpes-Côte d’Azur, Gis Ibisa, CNRS and Inserm.

## AUTHOR CONTRIBUTIONS

T.P. and G.J-M. contributed to metabolomic analyses; J-X.Y. and D.K performed transcriptomics analyses; M.E.M. performed the experiments and carried out data analyses; M.E.M. and M.G.F. conceived the study, designed the experiments, and wrote the manuscript. M.G.F. supervised the work.

## COMPETING INTERESTS

The authors declare that they have no competing interests.

## SUPPLEMENTARY FIGURES

**Supplementary figure 1:**
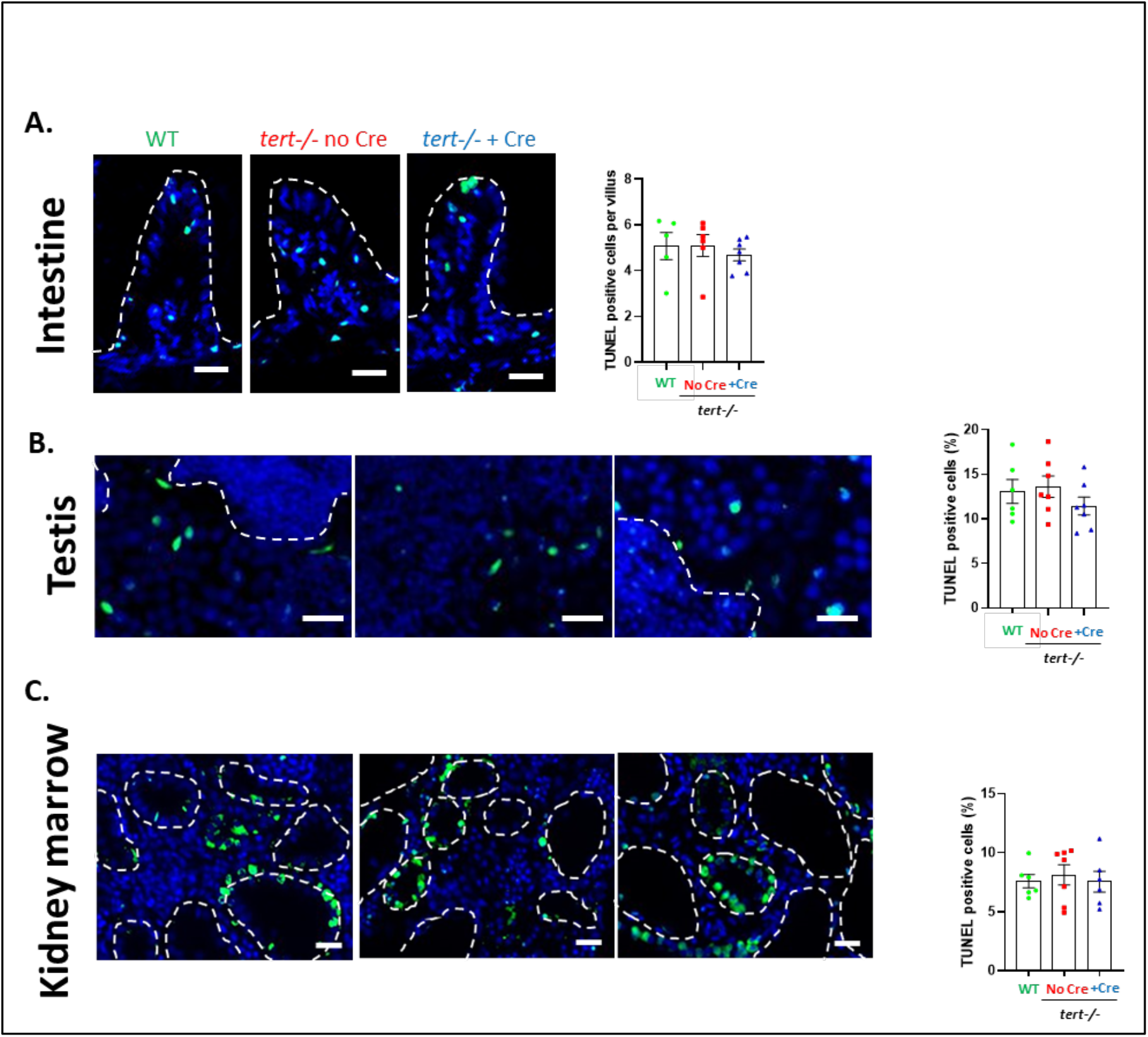
9-month-old fish did not exhibit apoptosis differences between conditions. **A-C**. Representative immunofluorescence images of apoptotic cell staining (TUNEL assay; left panel) and quantification (right panel) in gut (**A**.), testis (**B**.), or KM (**C**.) tissues of 9-month-old zebrafish. Scale bar: 20µm. Dashed lines delineate gut villi (**A**.), mature spermatid area (**B**.), or kidney tubules (**C**.). At 9 months of age, no differences in apoptosis were detected in gut, testis and KM comparing *tert-/-* No Cre, *tert-/-* +Cre and WT fish All data are represented as mean +/-SEM (N=5-7 per condition; no significance was detected comparing all conditions and using one-way ANOVA and post-hoc Tuckey tests).

**Supplementary figure 2:**
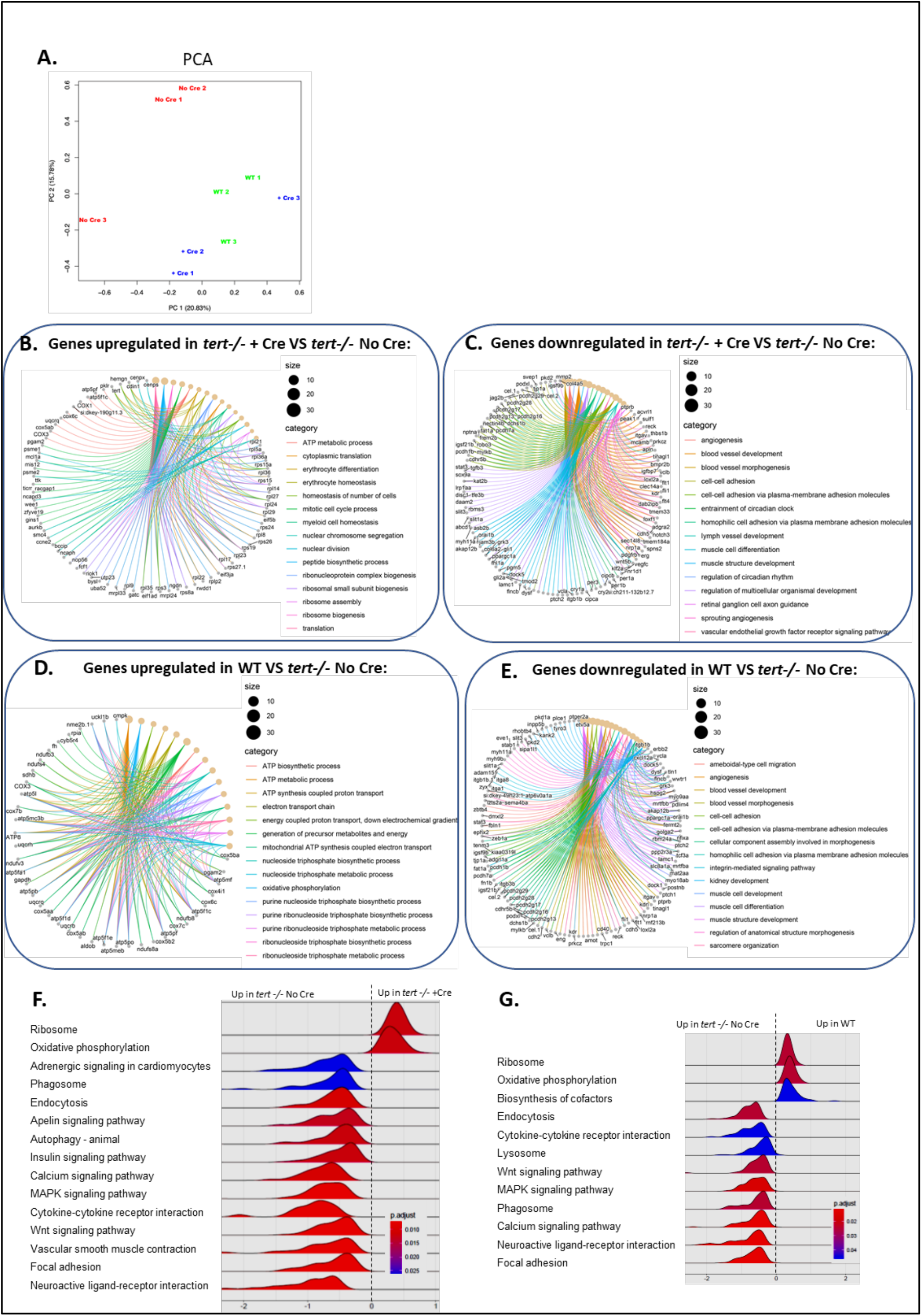
Gut specific *tert* expression rescues gut transcriptomic profile. **A**. Principal Component Analysis (PCA) based on untargeted transcriptomic data of 9-month-old gut samples. A clustering between *tert-/-* +Cre and WT while *tert-/-* No Cre group was clearly distinguishable from *tert-/-* No Cre fish (N=3 per group). **B-C**. Identification of Differentially Expressed Genes (DEGs) and their biological process categories (Gene Ontology -GO-Term analysis; FDR<0.1) that are downregulated **(B.)** or upregulated **(C.)** in the gut of 9-month-old *tert-/-* +Cre fish compared to *tert-/-* No Cre fish. **D-E**. Identification of Differentially Expressed Genes (DEGs) and their biological process categories (GO Term analysis; FDR<0.1) that are downregulated **(D.)** or upregulated **(E.)** in the gut of 9-month-old WT fish compared to *tert-/-* No Cre fish. Genes associated with morphogenesis, angiogenesis, cell-cell adhesion and muscle development are concomitantly downregulated compared to *tert-/-* No Cre reflecting the requirement of tissue repair in *tert-/-* No Cre fish. In parallel, ATP metabolism, ribonucleotide biosynthesis and mitotic cell cycle processes pathways are enriched in *tert-/-* +Cre and WT fish compared to *tert-/-* No Cre suggesting mitochondrial defects and reduced proliferation in *tert-/-* No Cre fish. **F-G**. Identification of KEGG (Kyoto Encyclopedia of Genes and Genomes) term using GSEA (Gene Set Enrichment Analysis) in the gut of 9-month-old tert-/-+Cre **(F.)** or WT **(G.)** fish compared to *tert-/-* No Cre. Genes associated with Ribosome and Oxidative phosphorylation pathways are enriched in the gut of both *tert-/-* +Cre and WT fish reflecting higher translation process and mitochondrial function compared to *tert-/-* No Cre. KEGG terms related to Cytokine-cytokine receptor interaction, Phagosome, Endocytosis, Neuroactive ligand-receptor interaction, Focal adhesion, Wnt signaling, MAPK signaling pathway and Calcium signaling pathway are reduced in both *tert-/-* +Cre and WT fish compared to *tert-/-* No Cre suggesting higher inflammation and requirement for tissue repair mechanisms.

**Supplementary figure 3:**
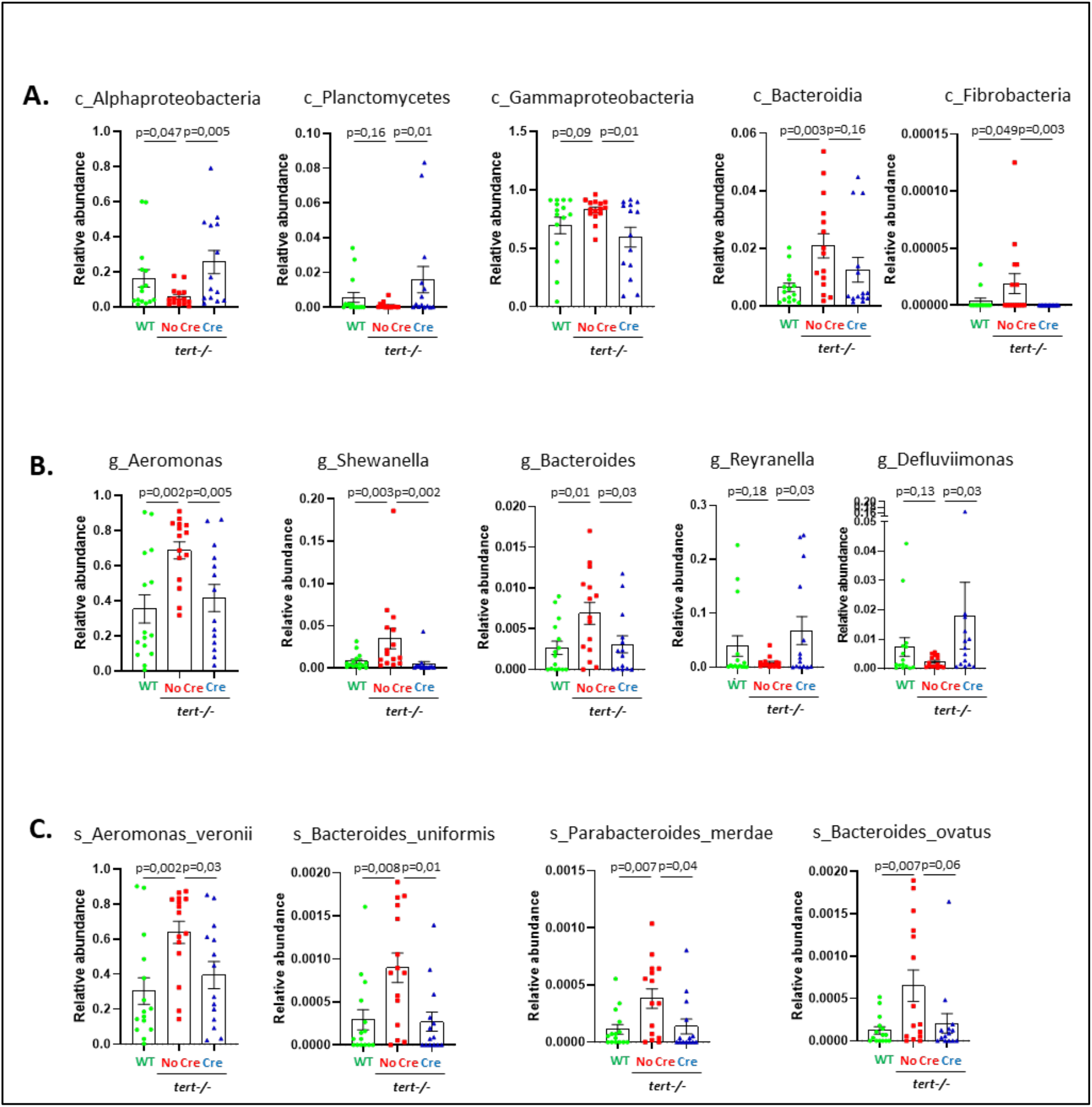
Gut-specific telomerase activity rescues alterations of gut microbiota composition. **A-C**. Relative abundance analysis of bacteria at the level of class **(A.)**; genus **(B.)** and species **(C.)**. N=14-15; p values were determined using Multiple hypothesis-test for sparsely-sampled features and false discovery rate (FDR). Tert expression in gut of tert-/-fish (*tert-/-* +Cre) recapitulates bacteria abundance at the class and species levels to WT profile compared to *tert-/-* No Cre where pathogenic bacteria are enriched.

**Supplementary figure 4:**
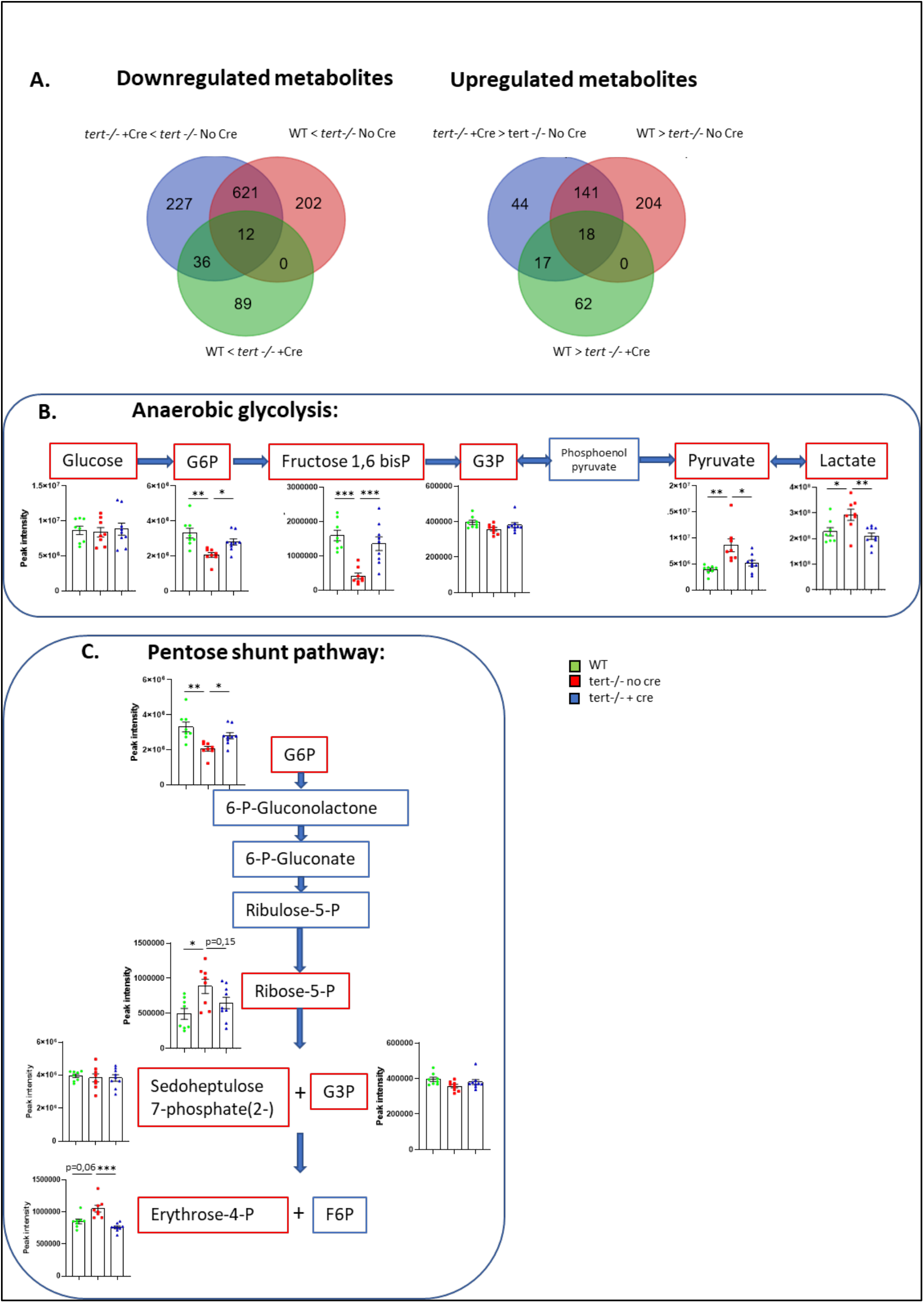
Gut-specific telomerase activity rescues gut metabolomic profile. **A**. Venn diagram representing downregulated (left panel) or upregulated (right panel) metabolites comparing the three conditions in gut of 9-month-old fish. The majority of metabolites detected in the gut of 9 months-old fish are concomitantly down or up-regulated in *tert-/-* +Cre and WT groups compared to *tert-/-* No Cre fish. **B-C**. Metabolomic analysis of the anaerobic glycolysis (**B**.) and pentose shunt pathways (**C**.) in gut of 9-month-old fish. Anaerobic glycolysis and pentose shunt metabolic profiles are rescued to WT levels in the gut of *tert-/-* +Cre compared *tert-/-* No Cre fish. All data are represented as mean +/-SEM (N=8-9 per condition; ^*^ p-value<0.05; ^**^ p-value<0.01, ^***^ p-value<0.001, using one-way ANOVA and post-hoc Tuckey tests). Red squares: detected metabolites; blue squares: undetected metabolites.

**Supplementary figure 5:**
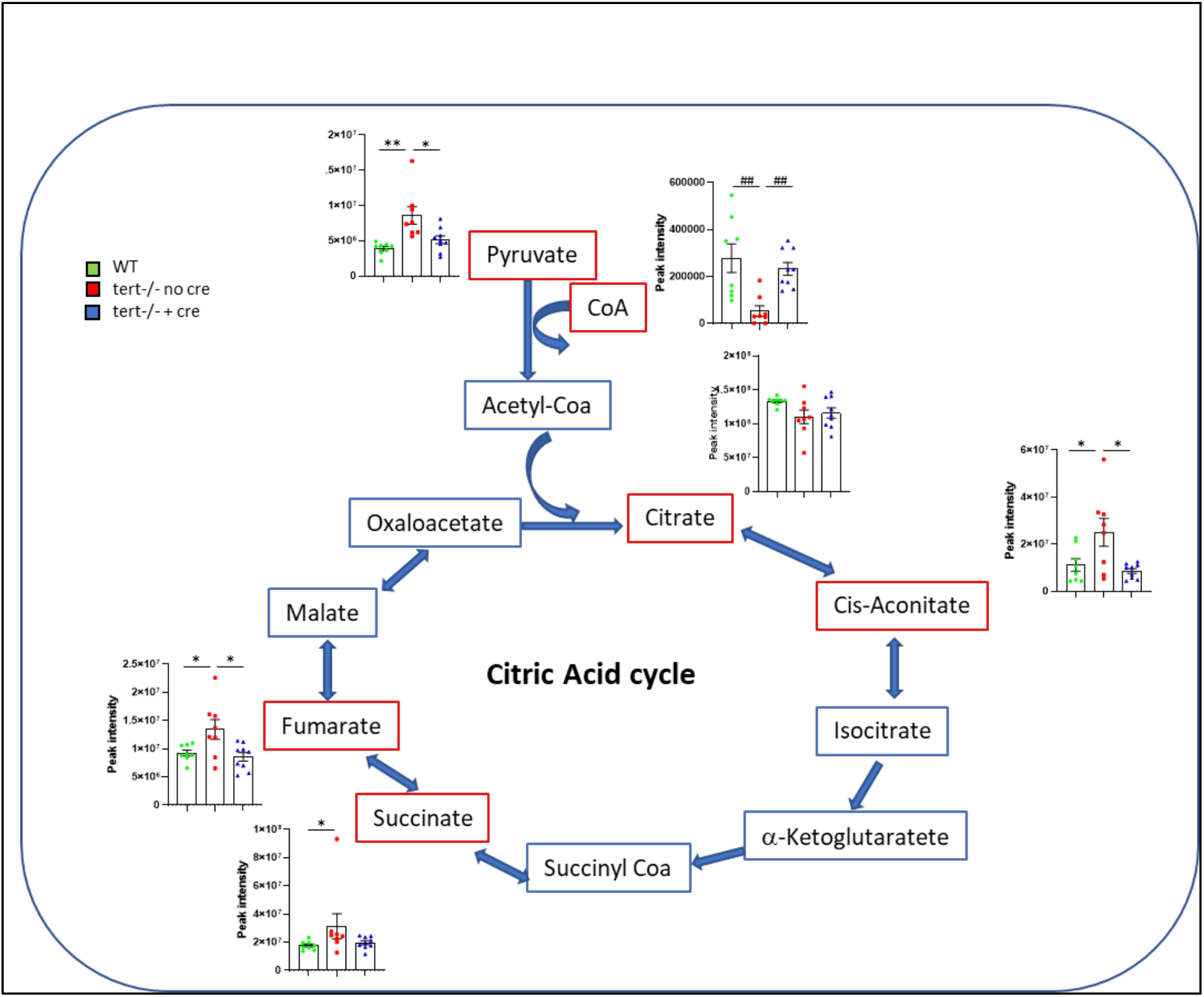
Gut-specific telomerase activity rescues citric acid cycle metabolism alterations in the gut of *tert-/-* fish. Metabolomic analysis of the citric acid cycle gut of 9-month-old fish. Citric cycle metabolic profile in the gut of *tert-/-* +Cre is similar to WT compared *tert-/-* No Cre fish. All data are represented as mean +/-SEM (N=8-9 per condition; ^*^ p-value<0.05; ^**^ p-value<0.01, using one-way ANOVA and post-hoc Tuckey tests; ## p-value<0.01, using Kruskal-Wallis and post-hoc Dunn’s tests). Red squares: detected metabolites; blue squares: undetected metabolites.

**Supplementary figure 6:**
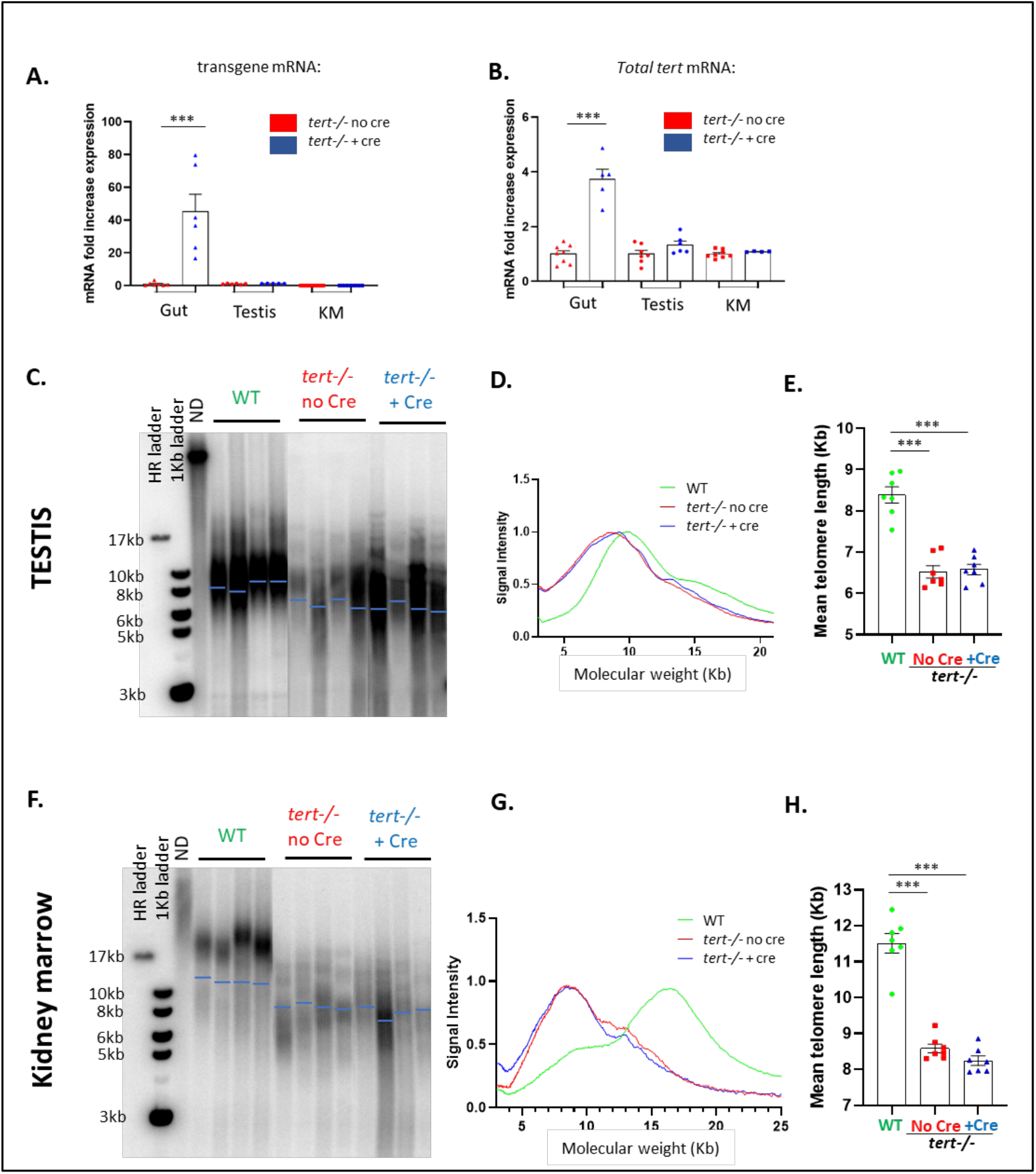
Cre-mediated *tert* transgene expression is specific to gut tissue. **A-B**. RT-qPCR analysis of *tert* transgene mRNA (**A**.) and total *tert* mRNA (**B**.; endogenous + transgene) expression in gut, testis of KM extracts from 9-month-old *tert-/-* No Cre and *tert-/-* +Cre fish. RT-qPCR graphs are representing mean ± SEM mRNA fold increase after normalization by *rps11* gene expression levels (N=5-8; ^***^ p-value<0.001, using one-way ANOVA and post-hoc Tuckey tests). While tert transgene transcription is induced by Cre injection in the gut of *tert-/-* fish compared to *tert-/-* No Cre, no transgene expression was detected in testis and KM of *tert-/-* +Cre fish. **C**. Representative images of telomere restriction fragment (TRF) analysis by Southern Blot of genomic DNA extracted from 9-month-old testis samples and quantifications of mean telomere length (blue bars). **D**. TRF mean densitometry curves from 9-month-old testis samples (N= 6-7). **E**. Quantification of mean telomere length analyzed by TRF on testis from 9-month-old fish. Data are represented as mean +/-SEM (N=6-7; ^***^ p-value<0.001, using one-way ANOVA and post-hoc Tuckey tests). **F**. Representative images of telomere restriction fragment (TRF) analysis by Southern Blot of genomic DNA extracted from 9-month-old KM samples and quantifications of mean telomere length (blue bars). **G**. TRF mean densitometry curves from 9-month-old KM samples (N= 6-7). **H**. Quantification of mean telomere length analyzed by TRF on KM from 9-month-old fish. Data are represented as mean +/-SEM (N=6-7; ^***^ p-value<0.001, using one-way ANOVA and post-hoc Tuckey tests). In accordance with lack of transgene expression in testis and KM, no difference in telomere length was detected in these organs when comparing *tert-/-* +Cre and *tert-/-* No Cre fish.

## Notes

### Competing Interest Statement

The authors have declared no competing interest.

